# Development of a Two-Leaf Photosynthetic Model Sensitive to Chlorophyll and Its Coupling with a Wheat Growth Model

**DOI:** 10.1101/2024.03.28.587183

**Authors:** Yining Tang, Jiacheng Wang, Yifan Hong, Caili Guo, Hengbiao Zheng, Xia Yao, Tao Cheng, Yan Zhu, Weixing Cao, Yongchao Tian

**Author notes:** E-mail addresses (Yining Tang);. edu.cn (Jiacheng Wang); (Yifan Hong); clguo@njau. edu.cn (Caili Guo); (Hengbiao Zheng); (Xia Yao); (Tao Cheng); (Yan Zhu); cao (Weixing Cao).

## Abstract

Crop growth models (CGMs) commonly simulates the response of photosynthetic rate to nitrogen (N) dynamic by calculating critical N concentration. However, critical N concentration makes it hard to describe the physiological effect of N dynamic on photosynthesis. Meanwhile, the effect of diffuse light on photosynthesis was limited in previous studies. In this study, we introduced a Two-leaf Photosynthetic Model Sensitive to Chlorophyll Content (TPMSCC) and coupled it with the crop growth model (WheatGrow) to enhance our understanding of how N dynamic and diffuse light affects photosynthesis. By coupling the Farquhar-von Caemmerer-Berry (FvCB) C_3_ photosynthesis model with a canopy radiative transfer model (PROSAIL), TPMSCC simulated the light interception of direct and diffuse light, and employed leaf chlorophyll content (LCC) to simulate the light absorption and electron transfer rate of leaves. Result showed that TPMSCC well simulated the light absorption of the wheat canopy and found the canopy photosynthetic rate benefited from the increase of diffuse radiation fraction (DRF) except for when the condition of a dense canopy at a high solar zenith angle was present. Research found the logarithmic and linear relationships of LCC to the initial light use efficiency (ɑ) and the maximum photosynthetic rate (A_max_), respectively, which followed the field measurements. Additionally, the optimized WheatGrow model outperforms its predecessor in describing the response of N application rate on photosynthesis.

## 1. Introduction

Crop growth models (CGMs) play a pivotal role in simulating crop growth, predicting yield, and assessing the effects of climate change and cultivation practices on yield formation (Dueri et al., 2022; Liu et al., 2019; Tang et al., 2023; Ye et al., 2020). However, CGMs may not adequately predict plant phenotype as consequence of genotype × environment × management (G × E × M) interactions for many key adaptive traits. For example, many CGMs rely on empirical functions to capture the relationship between phenotype and environmental factors, such as most models used nitrogen (N) stress factors to generalize the effect of N dynamics on photosynthetic assimilation (Manschadi et al., 2022; Soufizadeh et al., 2018; Zhao et al., 2014), neglecting the basis of photosynthetic physiology. This structured simplification of complex physiological processes was carried out to enable the application of CGM on large scale, assessing the effects of climate change, sowing date, and N application on crop yield (Liu et al., 2019; Nelson et al., 2002; Puntel et al., 2016; Ye et al., 2020). However, it is difficult to describe the phenotypic performance of multiple traits regulated by the interactions of G×E×M in crop breeding research (Cooper et al., 2014; Cooper and Hammer, 1996; Messina et al., 2011; Parent and Tardieu, 2014). Therefore, to address this limitation, there is a need to describe the response of phenotype to G×E×M by modeling physiological processes in crops, instead of developing the simple relationships between the phenotype and G×E×M (Soufizadeh et al., 2018).

Photosynthesis is an essential functional module in CGMs, including the simulation of light absorption and CO_2_ assimilation. CGMs commonly employed the Beer-Lambert law to model light interception, wherein incident light is considered direct and the canopy cover fraction (Liu et al., 2021) at solar direction is regraded as the fraction of photosynthetically active radiation (PAR; see Table 1 for the full explanation of all model variables) interception by the canopy (FIPAR). The extinction coefficient (k) calculated by leaf angle and solar zenith angle is used to simulate the leaf area projection at different heights within the plant canopy, so it describes the FIPAR at different canopy heights. FIPAR equals the fraction of PAR absorption by canopy (FAPAR) under the hypothesis of the full absorption of PAR by plant leaves and soil background (Widlowski, 2010). Some researchers distinguished the distribution of direct light and diffuse light in the plant canopy due to the various attenuation rates of diffuse light and direct light in the plant canopy. For instance, Boote and Pickering, (1994) used an empirical extinction coefficient to describe the attenuation of diffuse light. The above methods that did not distinguish the light absorption of sunlit leaves and shaded leaves were noted as a big-leaf model.

**Table 1:**
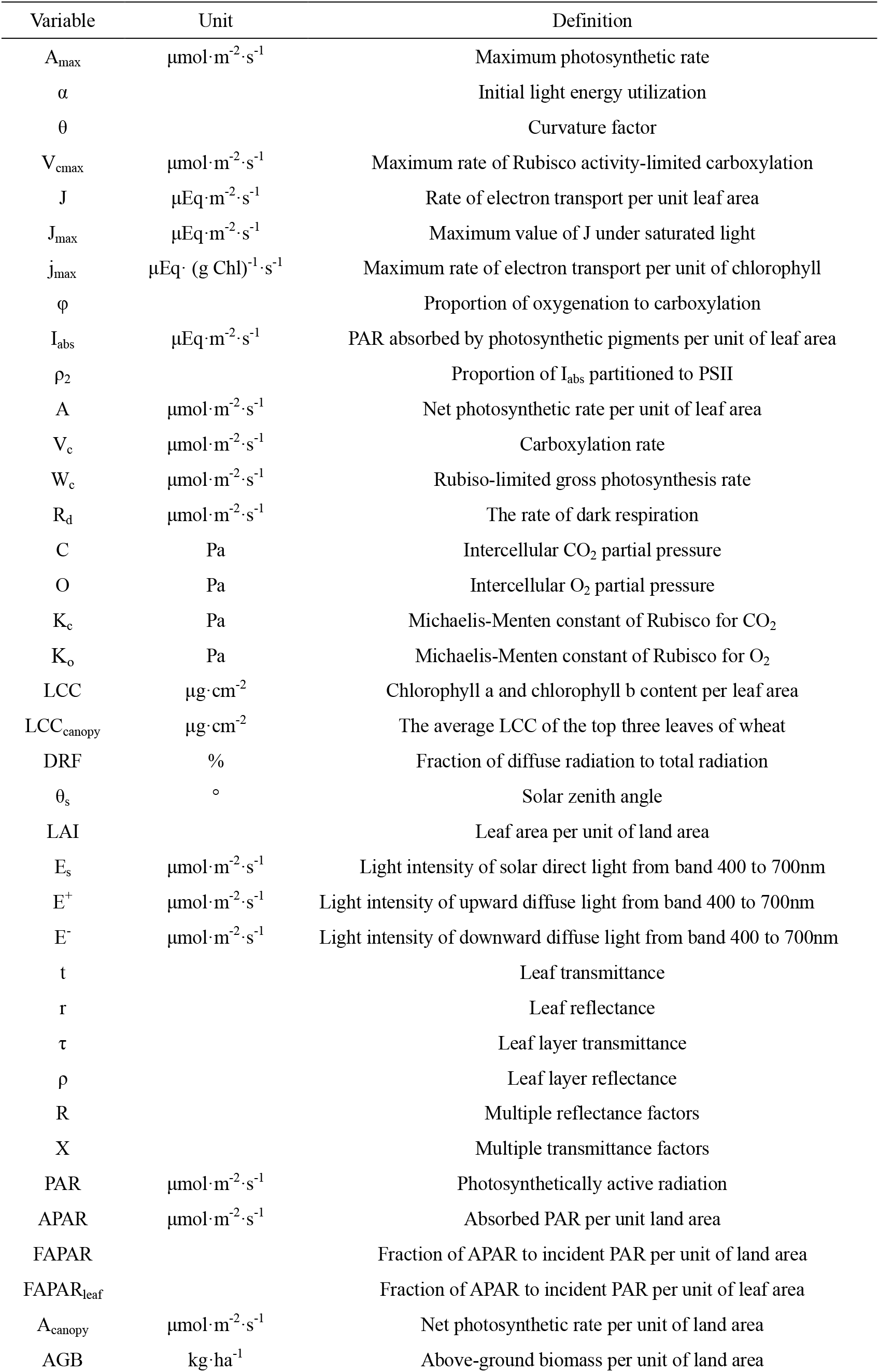
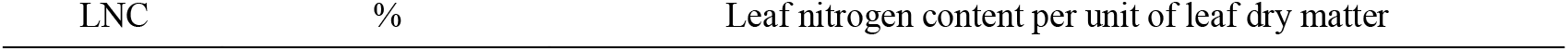
Full names of the abbreviations used in this study.

As the integral leaf area index (LAI) at canopy height increases, the distribution of light intensity within the canopy follows a pattern of exponential decline, consistent with the Beer-Lambert Law, but even the leaf at the bottom of the canopy may receive the transmitted high intense direct light. Thus, a two-leaf model (Boote and Pickering, 1994; Chen et al., 1999) distinguishing sunlit and shaded leaves was proposed. Shaded leaves are obstructed by upper canopy leaves, so the two-leaf model divided the plant canopy into several layers and estimated the fraction of shaded leaves in a specific layer by calculating the projected area of upper leaves in solar direction based on Beer-Lambert law, sunlit leaves absorb direct light and diffuse light, while shaded leaves absorb diffuse light.

However, Liu et al. (2021) indicated that commonly used CGMs have not adopted the two-leaf model but continue to rely on the big-leaf model. The lack of widespread adoption of the two-leaf model may partly be attributed to the absence of studies quantifying the biases introducing by big-leaf model. The application of CGM relies the model calibration using measured yield or biomass data (Huang et al., 2023; Kang and Özdoğan, 2019; Ye et al., 2020), which may offset and conceal the biases of photosynthesis in big-leaf model. Additionally, the diffuse light in canopy contains the direct light scattered by atmosphere and leaves, which is hard to simulate. Thus, there is no recognized method to describe the attenuation of diffuse light, existing studies use constant extinction coefficient to calculate the interception of diffuse light (de Wit et al., 2019; Shao et al., 2020), which dose not match reality. Hence, it is essential to develop a mechanistic model simulating diffuse light interception to explore the how the uncertainty of diffuse light fraction impacted photosynthesis.

CGMs often describe the impact of N dynamics on photosynthesis through the calculation of the N stress factor (de Wit et al., 2019; Jones et al., 2003). The factor represents the normalized actual N concentration within a range spanning from the minimum to critical N concentrations. Critical N concentration is the minimum N concentration necessary to achieve maximum above-ground biomass at any moment of vegetative growth (Lemaire et al., 1984). Photosynthetic parameters such as maximum photosynthetic rate (A_max_) and initial light energy utilization (α) are assumed to attain predetermined maximum value under critical N concentration (Manschadi et al., 2022; Soufizadeh et al., 2018; Zhao et al., 2014). This approach completely ignores the effects of elements like canopy structure and N allocation and attributes the phenomena of biomass gradually becoming saturated with higher N content to the saturation of photosynthetic parameters (Evans, 1989). The A_max_ and α need to be calibrated by the measured yield or biomass when the crop model was applied in a new ecological region based on the statistic method (Klosterhalfen et al., 2017; Ye et al., 2020), but the response of photosynthetic parameter to N dynamic has never been proved. Therefore, there is an urgent need for a mechanistic photosynthesis module that can improve the simulation of photosynthetic physiological processes and the response of N dynamics to dry matter accumulation.

In this study, we employed the Farquhar-von Caemmerer-Berry (FvCB) C_3_ photosynthesis model (Farquhar et al., 1980) to simulate the photosynthetic rate of single leaves. The leaf CO_2_ assimilation rate was set as the minimum value of the carboxylation rate restricted by the Rubisco enzyme (A_c_) or by electron transport (A_j_). A_c_ and electron transport rate (J) were the functions of the maximum carboxylation rate (V_cmax_) and maximum electron transport rate (J_max_), which were determined by the concentration of Rubisco enzyme and chlorophyll, respectively. There are several photosynthesis models based on FvCB model, which were applied in large-scale ecological studies, like the Boreal Ecosystem Productivity Simulator (BEPS, Chen et al., 1999) and Breathing Earth System Simulator (BESS, Jiang and Ryu, 2016). The response of photosynthesis to N in these models relies on empirical statistical models, such as the linear relationships between chlorophyll content, vegetation reflection spectrum, leaf N content, and V_cmax_ (Alton, 2017; Dechant et al., 2017; Luo et al., 2019). However, Braune et al. (2009) found that N concentration had sensitivity on initial light efficiency (ɑ) and dark respiration rate (R_d_). Thus, the effect of N on leaf photosynthesis cannot be fully explained if focusing only on V_cmax_. In addition, plants adjust the allocation of N to different leaf components (such as chlorophyll) according to the light environment, as to balance A_c_ and A_j_ to achieve an optimal photosynthetic carbon assimilation ability, namely the optimization theory (Chen et al., 1993; Evans, 1989; Wang et al., 2017). It indicates that, even with the same leaf N concentration, the same crop variety may have varied photosynthetic capacities due to varying light conditions. For example, plants will increase N allocation to chlorophyll to obtain higher J_max_, and reduce the synthesis of carboxylase enzyme under low illumination conditions, thus resulting in lower V_cmax_ (Evans, 1989). This indicates that the distribution of leaf N among the various photosynthetic components fluctuates depending on the environment, and therefore the estimation of the leaf photosynthetic parameter through total leaf N has some degree of inaccuracy. While chlorophyll content may be a more reliable photosynthetic indicator (Croft et al., 2017; Luo et al., 2019, 2018). FvCB can estimate the electron transfer rate per unit of leaf area by calculating the electron transfer rate of chlorophyll (Farquhar et al., 1980). A previous study indicated that the impact of N concentration on the conversion efficiency of incident light into J at the strictly limiting light level (k_2(LL)_) attributed to the variation of absorptance under different leaf chlorophyll content (LCC, Yin et al., 2009). The above information provides a theoretical framework correlating leaf chlorophyll content and photosynthetic rate.

Radiative transfer models were used for simulating the scatter characteristics of plant leaves and canopy. The leaf radiative transfer model PROSPECT (Jacquemoud and Baret, 1990) could simulate the response of leaf reflectance and transmittance on leaf biochemical components (e.g., chlorophyll, leaf dry matter). The canopy radiative transfer model SAIL (Verhoef, 1984) simulated the distribution characteristic of direct light, sky diffuse light, and the scattered light scattered by leaves and background. PROSAIL, the combination of PROSPECT and SAIL model, was widely used to simulate reflectance of canopy and retrieve features of plant canopy components (Dong et al., 2019; Jay et al., 2017; Jiang et al., 2020; Li et al., 2018; Verrelst et al., 2015; Zhao et al., 2017; Zhou et al., 2014), but was never used for photosynthetic simulation. Therefore, the effect of N dynamic on photosynthesis might be described from the aspects of photochemical and light absorption by a couple of PROSAIL and FvCB.

Thus, this study aims to: (1) develop a novel two-leaf photosynthetic model sensitive to LCC and diffuse radiation fraction (DRF); (2) investigate the response of LCC and DRF on light absorption and photosynthesis using the two-leaf photosynthetic model; (3) integrate the two-leaf photosynthetic model with the CGM (WheatGrow) and assess its performance.

## 2. Data and methodology

The data utilization and validation of this study are shown in Figure 1. This study used a modified PROSAIL model to simulate the vertical distribution of PAR in the wheat canopy, and the simulated FAPAR was validated using field measurements. Then, the PAR absorbed by photosynthetic pigments per unit of leaf area (*I_abs_*) was simulated by the PROSPECT model. *I_abs_*was put into the FvCB model to simulate leaf photosynthetic rate, A_max_ and α were extracted and validated by field measurements. Canopy photosynthetic rate was simulated by accumulating leaf photosynthetic rate of the whole canopy. Finally, the coupling of PROSAIL model and FvCB model, known as Two-leaf Photosynthetic Model Sensitive to Chlorophyll Content (TPMSCC), was incorporated into the WheatGrow model, resulting in WheatGrow-T. The field measurements were used to validate the simulated agronomic parameters of WheatGrow-T model.

**Fig. 1:**
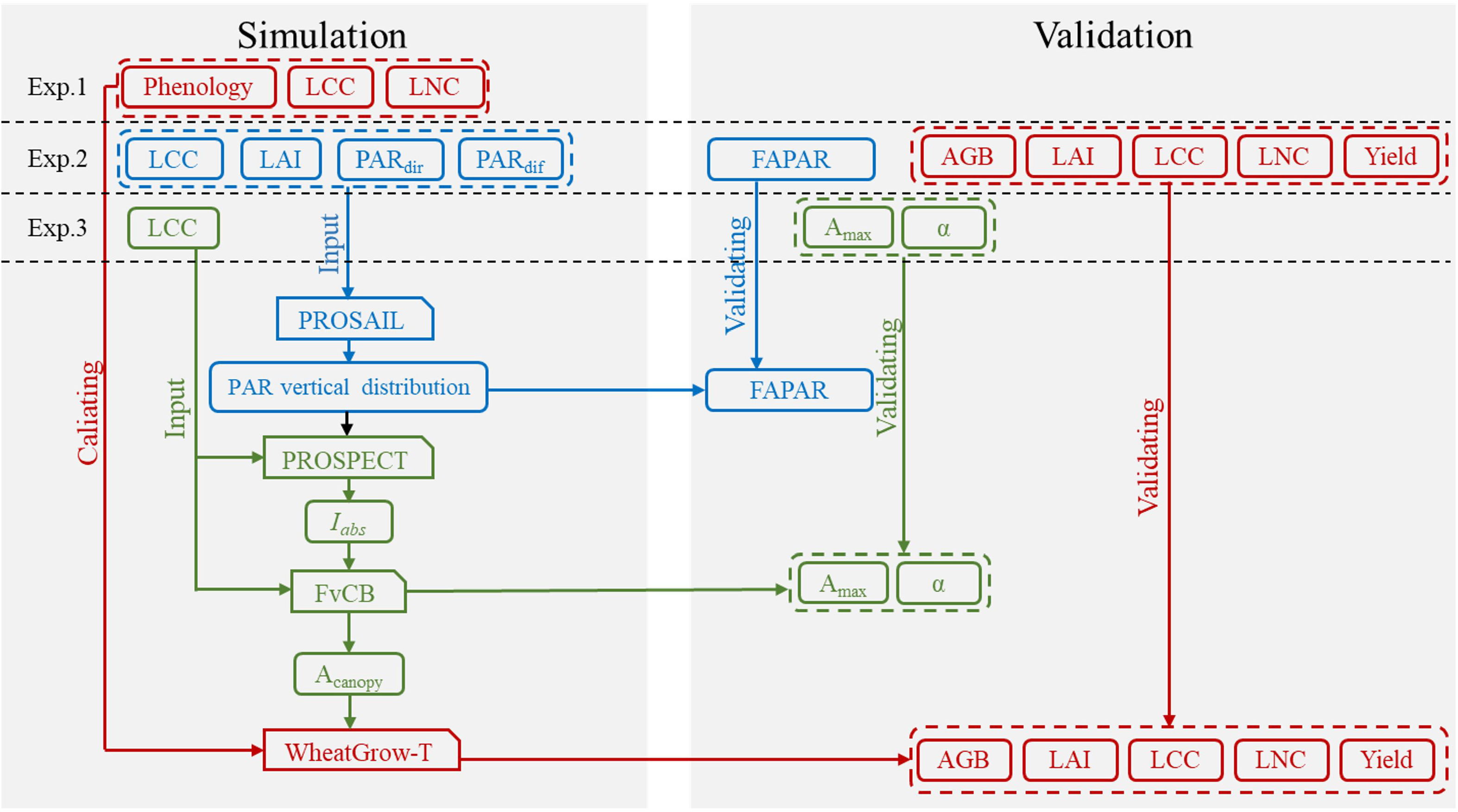
Overview of data utilization and validation.

### 2.1 Experimental design

The experimental area is located at the experimental demonstration base station of the National Engineering and Technology Center for Information Agriculture (NETCIA), Rugao, China (120°45′E, 32°16′N) from 2018 to 2020. Rugao County has an average annual temperature of 14.6 ℃, an average annual rainfall of 1055.5 mm, and its main soil texture is sandy soil and belongs to the main winter wheat production area in China. Three field experiments were conducted in this study, including treatments with different varieties and different N levels (Table 2).

**Table 2:**
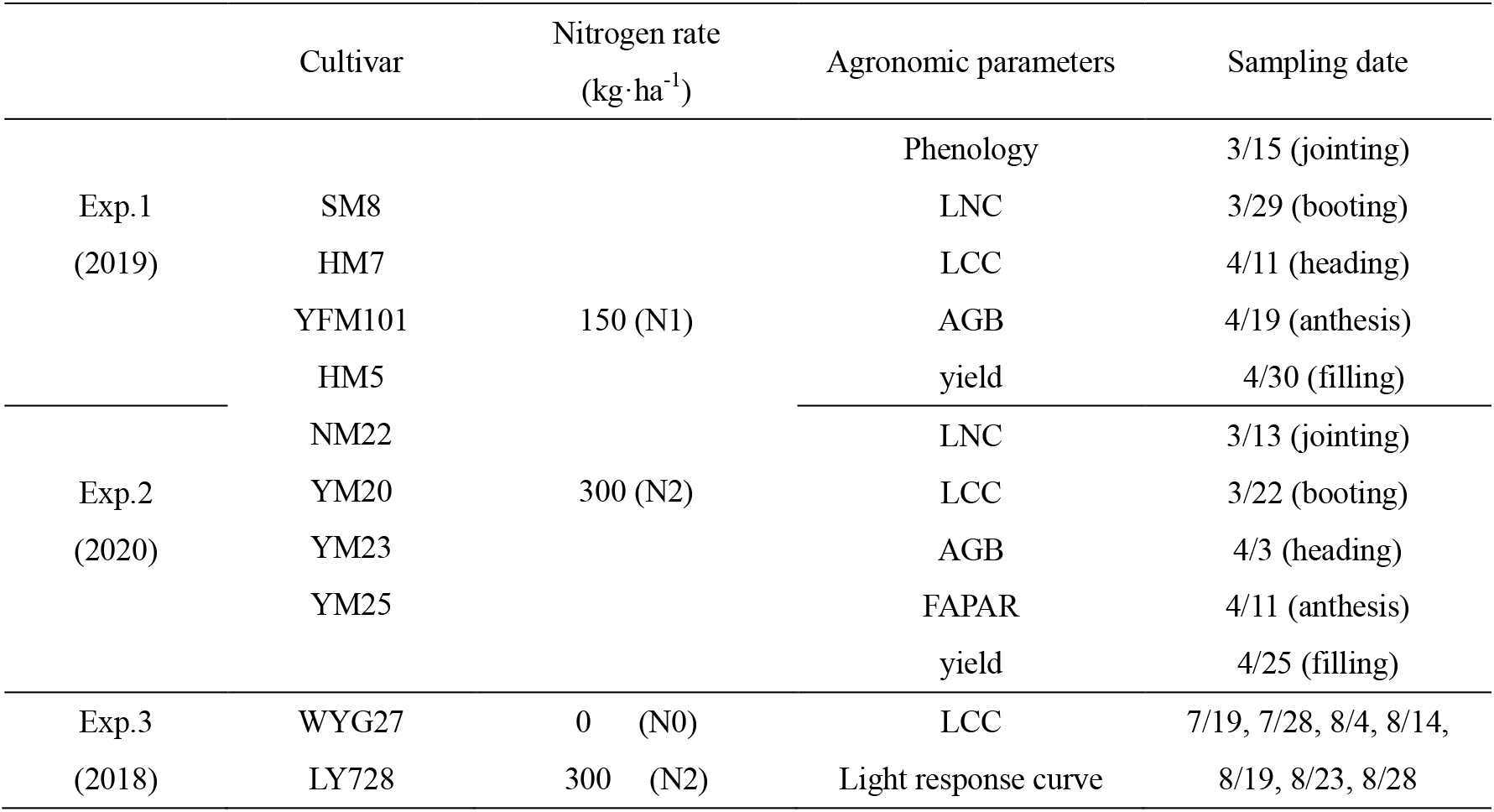
Experimental design and data acquisition.

Experiment 1 (Exp.1) was conducted from October 28, 2018 to May 29, 2019. Eight wheat varieties (Table 2) were sown on October 28, 2018, with a sowing amount of 150 kg·ha^-1^. Two N rates (N1: pure N 150 kg·ha^-1^, N2: pure N 300 kg·ha^-1^) were set up, half of the N fertilizer was applied before sowing and the remaining half was applied during the tillering stage. A phosphate fertilizer supplement of 420 kg·ha^-1^ P_2_O_5_ and a potash fertilizer of 120 kg·ha^-1^ K_2_O were applied to the experimental plots before sowing.

Experiment 2 (Exp.2) was conducted from November 2, 2019 to May 25, 2020. The wheat varieties, sowing amount, and fertilizer application were the same as Exp.1. Sowing date was November 2, 2019.

Experiment 3 (Exp.3) was conducted from June 18, 2017 to October 20. Two rice varieties (Table 2) and two N rates (N0: pure N 0 kg·ha^-1^, N2: pure N 300 kg·ha^-1^) were set, a total of four treatments were in Exp.3. Details of the Exp.3 can refer to Zhang et al. (2020).

### 2.2 Data acquisition

#### 2.2.1 Agronomic parameters

The data and acquisition period obtained in this study are shown in Table 2. The dates of booting, heading, and maturity stages of each treatment in Exp. 1 were documented. 30 plants were chosen to separate the stem, leaf, and spike from each wheat plant at each sampling point. The samples were withered at 105℃ for 20 min, dried at 80℃, and weighed to obtain the above-ground biomass (AGB). Leaf N concentration (LNC, %) was determined by vario MACRO cube (Elementar, Hanau, Germany). LAI was measured by LI-3000C Portable Leaf Area Meter (Li-Cor, Lincoln, Nebraska, USA). The relative chlorophyll content of the top three leaves in 10 plants was determined by Dualex 4 Scientific leaves meter (Force-A, Orsay, France), and then converted to true LCC (Xu et al., 2019). The average canopy leaf chlorophyll content (LCC_canopy_) was calculated by the average value of the top three leaves. At the maturity stage, one square meter of wheat was harvested and dried at 80℃, and the grain yield per unit of land area was recorded. Soil nutrient content was measured by executing a third party.

#### 2.2.2 Canopy radiation parameters

In Exp.2, the canopy analyzer SunScan (Delta-T, Cambridge, UK) was used to measure the PAR above the canopy (PAR_top_) and the PAR at the bottom of the canopy (PAR_bottom_) at jointing, booting, and heading stages. In addition, the diffuse PAR (PAR_dif_) and the direct PAR (PAR_dir_) above the canopy were measured by the diffuse reflection coefficient sensor BFS. FIPAR was calculated based on equation (1). The previous study (Widlowski, 2010) found a slight difference between FAPAR and FIPAR under the situation of green leaves and black background, thus the FIPAR calculated here was noted as FAPAR in this study.

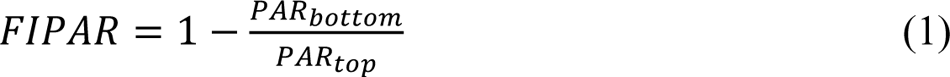

#### 2.2.3 Light response curve

In Exp.3, light response curves of blade for different canopy height were measured using a LICOR-6400XT Portable Photosynthesis System (LI-COR). PAR was set to 2000, 1750, 1500, 1200, 1000, 750, 500, 250, 100, 60, 20, and 0 μmol·m^-2^·s^-1^ to obtain light response curves, the net photosynthetic rates of various PARs were fitted. Based on the non-rectangular hyperbola (2, Prioul and Chartier, 1977), the maximum photosynthetic rate (*A_max_*), initial light energy utilization (α), and respiration rate (*R_d_*) of leaves were obtained.

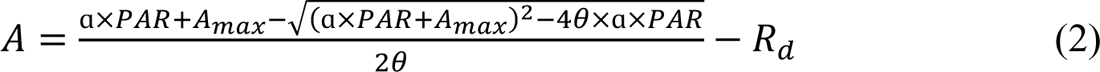

Where, *A* is the net photosynthetic rate (µmol·m^-2^·s^-1^); *PAR* is photosynthetically active radiation (µmol·m^-2^·s^-1^); *A_max_*is the maximum photosynthetic rate at light saturation (µmol·m^-2^·s^-1^); *α* is the initial slope of light response curve when PAR is 0-200 µmol·m^-2^·s^-1^; *R_d_* is dark respiration rate (µmol·m^-2^·s^-1^); *θ* is the curvature factor.

#### 2.2.4 Meteorological data

Meteorological information for Experiments 1 and 2 was gathered from the Rugao meteorological station. The weather information includes daily minimum and maximum temperatures, sunshine duration, and precipitation. Daily PAR_dir_ and PAR_dif_ were calculated according to sunshine hours (Spitters et al., 1986).

### 2.3 WheatGrow-T model

#### 2.3.1 Overview

A Two-leaf Photosynthetic Model Sensitive to Chlorophyll Content (TPMSCC) was proposed, which is a combination of PROSAIL and FvCB, the input parameters of TPMSCC are shown in Table 3. TPMSCC was integrated with WheatGrow to construct a novel crop model (WheatGrow-T, Fig.2), which is driven by chlorophyll. Meanwhile, a big-leaf photosynthetic model considered DRF and chlorophyll, named Big-leaf_sep_, was developed for comparison, its coupling with WheatGrow generated WheatGrow-B. The original WheatGrow was also used for comparison, the photosynthetic module named Big-leaf ignored DRF and chlorophyll (Table 4) and described the effects of N dynamic on photosynthetic parameters using critical N concentration.

**Fig. 2:**
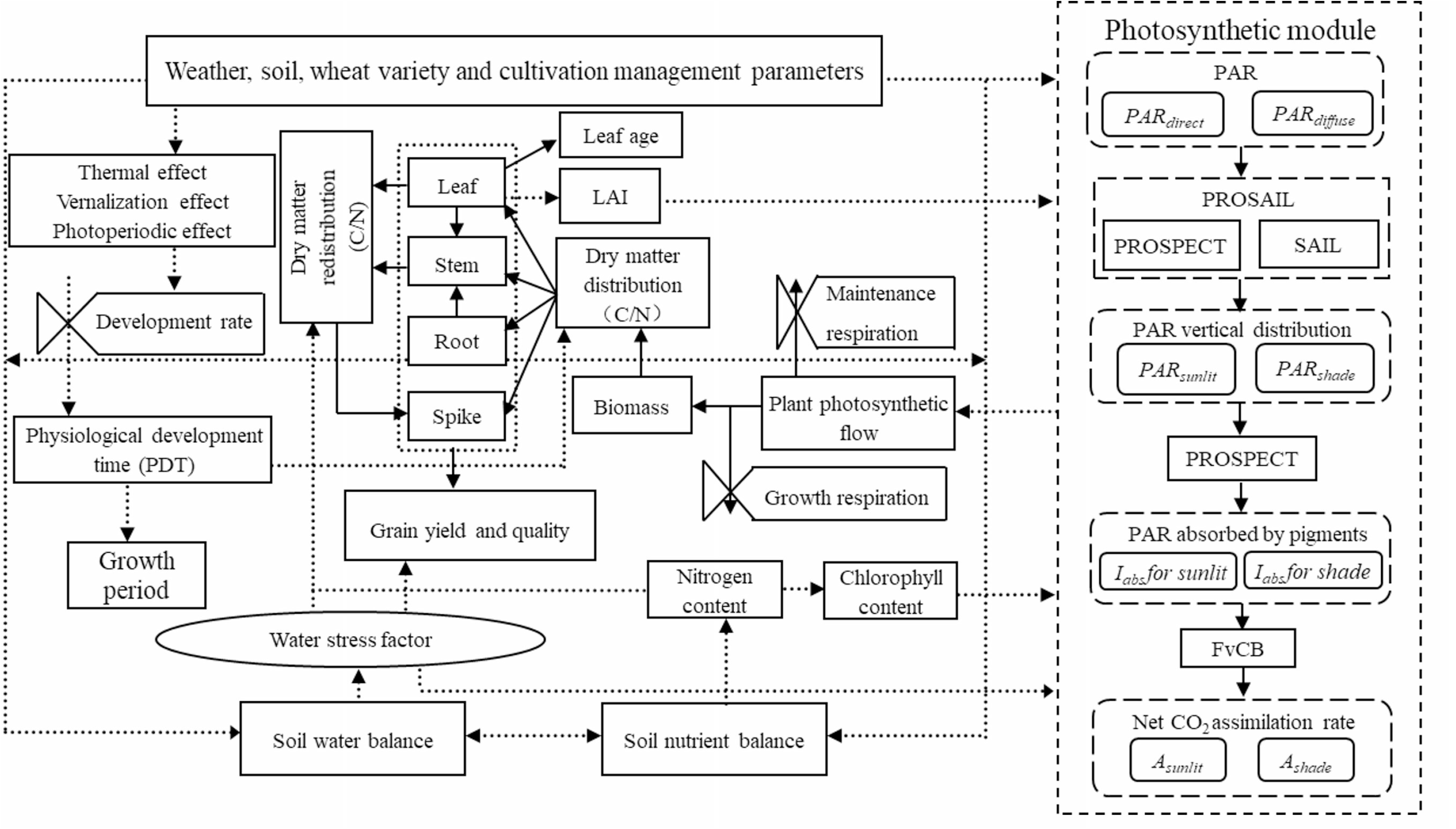
Modular structure of the WheatGrow-T model.

**Table 3:** Input parameters used in the TPMSCC model.

| Parameter | Abbr. in model | Unit |
| --- | --- | --- |
| **PROSPECT** |  |  |
| Carotenoid content | Car | μg·cm<sup>-2</sup> |
| Dry matter content | C<sub>m</sub> | g·cm<sup>-2</sup> |
| Leaf structural coefficient | N | — |
| Chlorophyll content | LCC | μg·cm<sup>-2</sup> |
| Brown pigment content | Cbrown | — |
| Equivalent water thickness | C<sub>w</sub> | cm |
| **SAIL** |  |  |
| Mean leaf inclination angle | θ<sub>l</sub> | ° |
| Leaf area index | LAI | — |
| Intensity of direct light | PAR<sub>dir</sub> | μmol·m<sup>-2</sup>·s<sup>-1</sup> |
| Intensity of diffuse light | PAR<sub>dif</sub> | μmol·m<sup>-2</sup>·s<sup>-1</sup> |
| Solar zenith angle | θ<sub>s</sub> | ° |
| **FvCB** |  |  |
| Chlorophyll content | LCC | μg·cm<sup>-2</sup> |
| PAR absorbed by pigments | I<sub>abs</sub> | μmol·m<sup>-2</sup>·s<sup>-1</sup> |

**Table 4:** Features of models developed in this study.

| **Table 4:** Features of models developed in this study |  |  |  |  |
| --- | --- | --- | --- | --- |
| Photosynthetic model | Consider chlorophyll | Distinguish PAR<sub>dir</sub> and PAR<sub>dif</sub> | Distinguish sunlit and shaded leaves | Crop model |
| TPMSCC | ✓ | ✓ | ✓ | WheatGrow-T |
| Big-leaf<sub>sep</sub> | ✓ | ✓ | ✗ | WheatGrow-B |
| Big-leaf | ✗ | ✗ | ✗ | WheatGrow |

#### 2.3.2 Description of TPMSCC

Photosynthesis is divided into light reaction and dark reaction. Light energy is firstly absorbed and is converted to electrical energy in the light reaction. In this process, antenna pigment-protein complexes containing chlorophyll *a+b* and the reaction center complexes containing chlorophyll *a* are involved in light absorption and electron transfer (Jin et al., 2016), respectively. Then, CO_2_ is reduced to carbohydrates in a dark reaction by the Rubisco enzyme, which was a theoretical basis for linking N dynamic and photosynthesis (Croft et al., 2017). The relationship between chlorophyll content and light absorption was simulated by the PROSAIL model; FvCB was used to calculate the electron transfer rate under given light absorption and CO_2_ assimilation rate.

##### 1. Radiation absorption simulation

The simulation of the vertical distribution of PAR and the absorption of PAR per unit of land area (APAR) was based on the modified PROSAIL model (Yang et al., 2017), which simulated scatter features of leaf and homogeneous canopy. In the PROSAIL model, direct light (E_s_), upward diffuse light (E^+^), downward diffuse light (E^-^), and radiance in the observer’s direction (E°) of the canopy are calculated by the following equations:

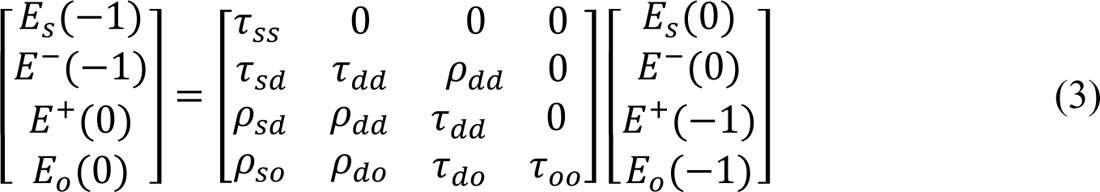

Where 0 and −1 behind E represent the top and bottom of the canopy, respectively; *τ* and *ρ* are transmittance and reflectance of the isolated canopy layer, respectively; the double subscripts indicate types of flux on incidence and exit, respectively. *s* stands for solar, *d* for diffuse hemispherical, and *o* for flux in the observer’s direction. *ρ* and *τ* were determined by leaf transmittance (*t*), reflectance (*r*), LAI, leaf angle, and solar position according to the SAIL model (Verhoef, 1984). *t* and *r* were determined by leaf components (such as LCC and carotenoid content) according to the PROSPECT model.

TPMSCC divided a homogeneous canopy into 60 heterogeneous leaf layers by numbers from 1 to 60 and simulated the distribution of *PAR_dir_*and *PAR_dif_* in each leaf layer. LCC of each leaf layer was calculated by the proportion of LCC in the top three leaves and LCC_canopy_ in Exp.1. The vertical distribution of LAI was calculated by the function as follows:

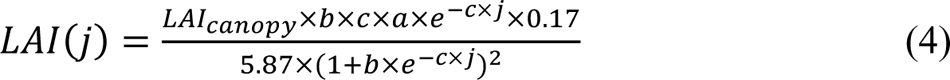

Where fitted parameters *a*, *b*, and *c* were set as 6.34, 25.51, and 6.27, respectively (Li et al., 2010); *LAI_canopy_* is integrated LAI of the canopy; *j* is relative height ranges from 1 to 60.

For leaf layer j = 60 to 1 (top to bottom):

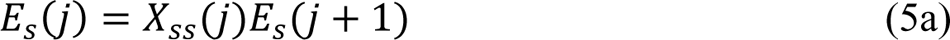

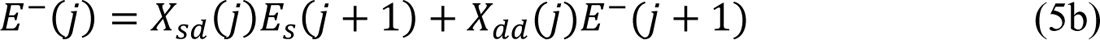

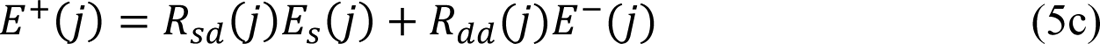

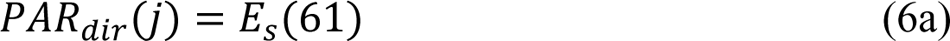

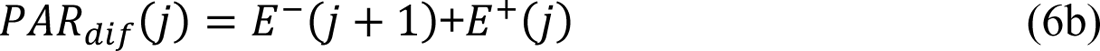

Where *E_s_(61)* and *E^-^(61)* are the incident solar direct light and sky diffuse light, respectively. *R* and *X* are multiple reflectance and transmittance factors (Yang et al., 2017), respectively, which were calculated from the bottom to top (j = 1 to 60):

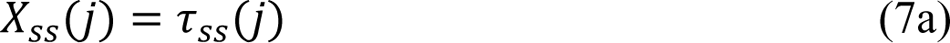

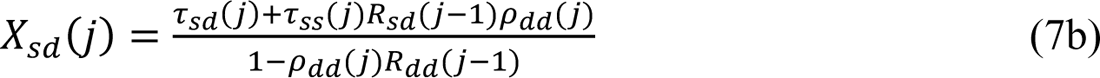

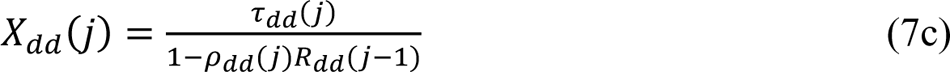

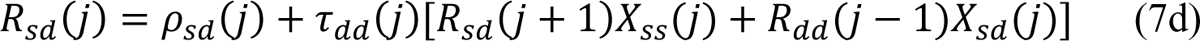

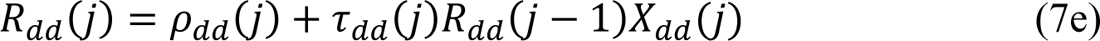

Where *R_sd_(0)* and *R_dd_(0)* are soil reflectance.

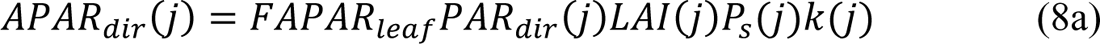

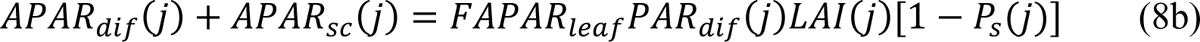

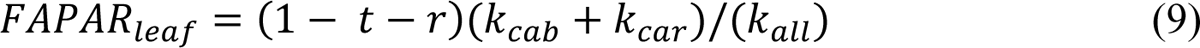

Where *APAR* is the absorption of PAR by photosynthetic pigments (chlorophyll and Carotenoid); *FAPAR_leaf_*is the fraction of PAR absorbed by photosynthetic pigments per unit of leaf area; *APAR_dir_* and *APAR_dif_* are APAR of solar direct light and sky diffuse light, while *APAR_sc_* is the APAR of diffuse light scattered from leaves and soil background; *k_cab_* and *k_car_* are absorption coefficients of chlorophyll and carotenoid, which are contained in PROSPECT model; *k_all_* is the sum of absorption coefficients of all leaf components in PROSPECT; *P_s_*is the fraction of sunlit leaves.

By setting soil reflectance as 0 (*R_sd_(0)* and *R_dd_(0)* equal 0), and also setting leaf reflectance and transmittance as 0 (*t* and *r* equal *0*) when computing *τ* and *ρ*, the outcome of equation (8b) exclusively encompasses *APAR_dif_*. This approach eliminates the contribution of scattered light from both leaves and soil, thereby allowing the differentiation between *APAR_dif_* and *APAR_sc_*, and analyzes the importance of considering *APAR_sc_*.

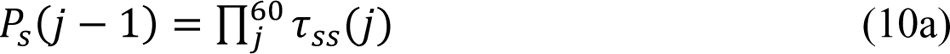

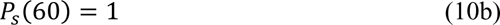

Sunlit leaves absorb both *PAR_dir_*, *PAR_dif_*, but shaded leaves absorb just *PAR_dif_*. Big-leaf_sep_ does not distinguish the sunlit and shaded leaves and allocates *PAR_dir_*to all leaves in each leaf layer. A description of Big-leaf can be found in the light interception of the APSIM model (Zheng et al., 2015), the variation of DRF and LCC were all ignored.

##### 2. Photosynthetic rate simulation

Leaf net photosynthetic rate was simulated by the FvCB model (Von Caemmerer et al., 2009), CO_2_ assimilation net rate of (*A*) was calculated as:

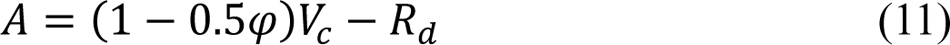

Where *V_c_* is the carboxylation rate (μmol·m^-2^·s^-1^); *φ* is the proportion of oxygenation to carboxylation, which was set as 0.27 for C_3_ plants (Farquhar et al., 1980); *R_d_* is the rate of dark respiration.

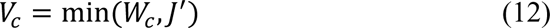

Where *W_c_*is the Rubiso-limited gross photosynthesis rate (μmol·m^-2^·s^-1^); *J’* is the RuBP-limited gross photosynthesis rate (μmol·m^-2^·s^-1^).

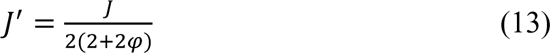

Where *J* is the rate of electron transport per unit of leaf area (μEq·m^-2^·s^-1^).

A non-rectangular hyperbola was used to describe the response of *J* to the light absorbed by the photosystem:

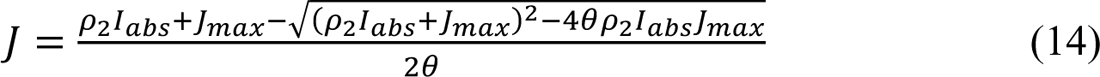

Where *J_max_* is the maximum rate of electron transport; *θ* is the curvature factor which was set as 0.8 in this paper; *I_abs_* is the PAR absorbed by leaf photosynthetic pigments, different from APAR, *I_abs_* is the light absorption per unit of leaf area; *ρ_2_*is the proportion of *I_abs_* partitioned to the photosystem II, and was set as 0.5 typically (Von Caemmerer et al., 2009; Yin et al., 2006).

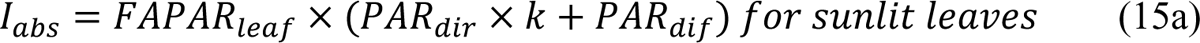

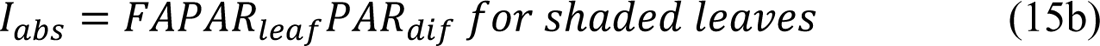

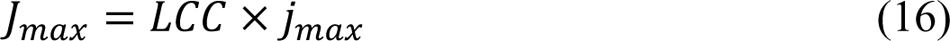

Where *j_max_* is the maximum rate of electron transport expressed on a chlorophyll basis as μEq· (g Chl)^-1^·s^-1^, and was set as 467 μEq· (g Chl)^-1^·s^-1^ when 25 ℃ (Farquhar et al., 1980).

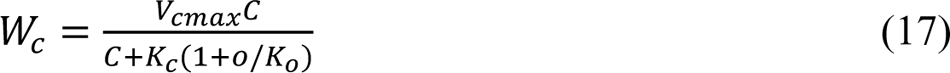

*V_cmax_* is the maximum velocities of the carboxylase at 25℃; *C* and *O* are the intercellular partial pressures of CO_2_ and O_2_, respectively, and are set as 26.24 Pa and 20500 Pa, respectively (Ryu et al., 2011); *K_c_* and *K_o_* are the Michaelis-Menten constants for CO_2_ and O_2_, respectively, and set as 40.4 Pa and 24800 Pa at 25℃, respectively (De Pury and Farquhar, 1997).

*V_cmax_* was calculated by *J_max_* according to (Wullschleger, 1993):

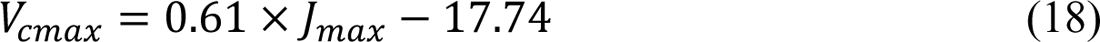

Archontoulis et al. (2012) found that RuBP-limited gross photosynthesis rate could describe the entire light response curve when *W_c_* is always higher than *J^’^*, and leaf net photosynthetic rate varied with PAR was expressed as:

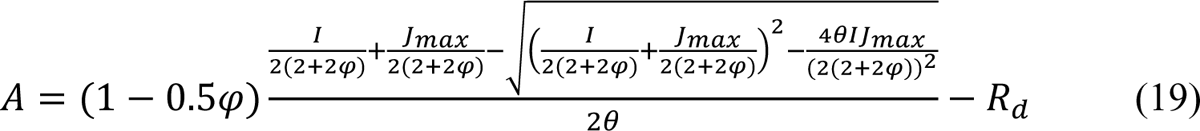

Reconstructing TPMSCC by the light response curve (2), getting the relationship of LCC to *A_max_*and *α*:

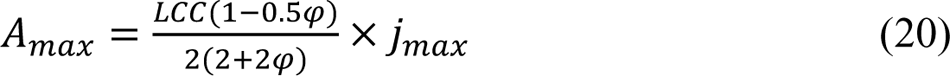

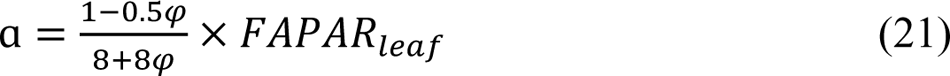

If the increase in light intensity leads to Rubisco-limited photosynthetic rate becoming the limiting factor of photosynthesis, *A_max_* is Rubisco-limited photosynthetic rate:

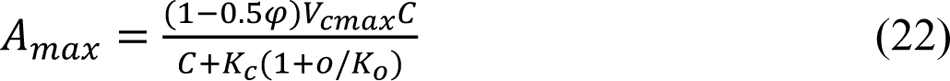

The net photosynthetic rate of each leaf layer (A_leaflayer_) is the sum of the net photosynthetic rate of sunlit leaves and shaded leaves:

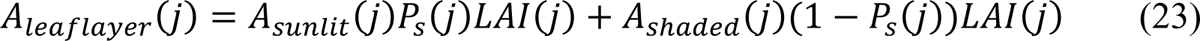

Canopy net photosynthetic rate (A_canopy_) is the sum of each leaf layer:

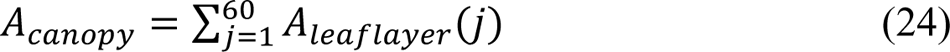

#### 2.3.3 Coupling of WheatGrow and TPMSCC

The WheatGrow-T model was constructed by replacing the photosynthetic module of WheatGrow with TPMSCC. Since chlorophyll was used as the intermediate variable linking N dynamic and photosynthesis, the N dynamic in the WheatGrow model that was originally based on critical N concentrations needed to be modified. A linear model based on the high correlation between LCC and LNC was used to transform the LNC simulated by WheatGrow to LCC. (Cheng et al., 2018; Schlemmer et al., 2013). The maximum LCC_canopy_ of specific wheat cultivar was stable annually (Fig. S1), thus the maximum LCC_canopy_ was set as a cultivar parameter, named Cab_max_. LCC_canopy_ was linearly related to LNC when LCC_canopy_ was lower than Cab_max_, resulting in the response of photosynthetic parameters to LNC. However, the increased LNC had no effects on photosynthetic parameters when LCC_canopy_ saturated. The minimum N concentration corresponded to the N concentration with no chlorophyll and was defined as the structural N concentration in previous studies.

### 2.4 Data utilization and analysis

#### 2.4.1 Model calibration

Parameters of crop models that required calibration were summarized in Table 5.

**Table 5:** Parameters in crop models requiring calibration.

| Parameter | Abbr. in model | Unit | Source |
| --- | --- | --- | --- |
| Intrinsic earliness | IE | — | Field data calibration |
| Photoperiod sensitivity | PS | — | Field data calibration |
| Filling duration factor | FDF | — | Field data calibration |
| *Parameters specific to WheatGrow-T and WheatGrow-B* |  |  |  |
| Maximum chlorophyll content | Cab<sub>max</sub> | µg·cm<sup>-2</sup> | Field data calibration |
| Conversion coefficient from LNC to LCC | Slope |  | Field data calibration |
| *Parameters specific to WheatGrow* |  |  |  |
| Critical N concentration | — | % | Field data calibration |
| Maximum photosynthetic rate | A<sub>max</sub> | µmol·m<sup>-2</sup>·s<sup>-1</sup> | Field data calibration |
| Initial light energy utilization | α |  | Field data calibration |

Field measurements of Exp.1 were used for model calibration.

The calibration of WheatGrow-T and WheatGrow-B was performed as follows: Firstly, phenological parameters: basic earliness, photoperiod sensitivity, and filling factor, were optimized by measured values of booting, heading, and maturity stages using particle swarm optimization algorithm (PSO, Xi et al., 2015); Then, Cab_max_ was set as the maximum value of LCC_canopy_ measured from each wheat cultivar; Finally, the conversion function from LNC to LCC_canopy_ was determined.

The conversion equations between LNC and LCC_canopy_ in different growth stages were developed. It was shown that leaves of early stages required a higher LNC than late stages to reach the same LCC_canopy_ (Fig. S2a). Assuming that the structural N content of wheat leaves was constant at different growth stages, leaf structural N content was set as 0.5 g·g^-1^, the measured minimum value of LNC in the bottom of the canopy at the filling stage. Thus, the linear equation of LNC and LCC_canopy_ at different growth stages went through the point of (0.5, 0), and obtained the linear equation of LCC_canopy_ at different growth stages (Fig. S2a). The slope of the equation was the allocation coefficient of leaf N to chlorophyll except for the structural N. By fitting the linear equation (Fig. S2b) of the allocation coefficient of different growth periods with physiological development time (PDT), the conversion equation of days after sowing was obtained.

The calibration of the WheatGrow model was performed as following: phenological parameters were firstly calibrated. Then, critical N concentration was calibrated, LNC and AGB of each wheat variety were employed to generate critical N dilution curves (Makowski et al., 2020), and the critical N concentration of each growth stage was extracted from critical N dilution curves (Zhao et al., 2014); Finally, A_max_ and α of each wheat cultivar were calibrated through measured AGB, two parameters were optimized to minimize the error between the simulated and measured AGB, the PSO was employed as optimization algorithm.

#### 2.4.2 Model evaluation

The sensitivity of photosynthetic rate to DRF was firstly validated with the measured FIPAR from Exp.2. Firstly, the incident PAR_dir_, PAR_dif_ above canopy measured by SunScan, and solar zenith angle were input into the TPMSCC and Big-leaf model to simulate the PAR value in the bottom of canopy, the simulated FAPAR calculated by equation (1) was recorded, RMSEs (25) between the simulated FAPAR and measured FAPAR were calculated. The vertical distribution of the simulated APAR was then subjected to a sensitivity analysis under various DRF, canopy densities, and sun zenith angles. Finally, the sensitivity analysis of simulated A_canopy_ under different PAR values (canopy light response curve) was conducted, and two scenarios were considered: one is natural and the other is ideal illumination. According to Spitters et al. (1986), the change rule for natural illumination is that DRF varies in response to PAR values; the ideal illumination was proposed in which DRF is consistent and independent of light intensity.

The sensitivity of photosynthesis to LCC was then validated with the measured A_max_ and α from Exp.3. Firstly, the relationship between LCC and the simulated A_max_ and α was compared to the measured results. Then, the measured LCC of each treatment was inputted in TPMSCC to simulate A_max_ and α. The simulation was validated by calculated RMSE and R^2^ (26) between the simulated and measured values. The effects of LCC on canopy light response curve were also analyzed.

Finally, the performance of WheatGrow-T was validated using the measured values from Exp.2. The simulated time sequences of AGB for WheatGrow-T, WheatGrow-B, and WheatGrow were compared with measured values. The sensitivity of N application rate to agronomy parameters (LNC, LCC, and AGB) under different Cab_max_ was conducted, which was based on both WheatGrow-T simulations and field measurements. The simulated results for LAI, LCC, LNC, AGB, and yield were verified against the measured values. R^2^ and RMSE values were used for evaluation.

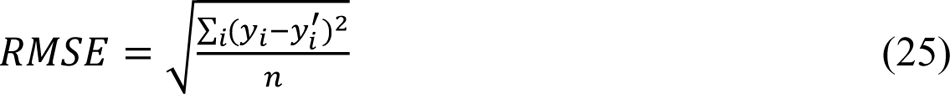

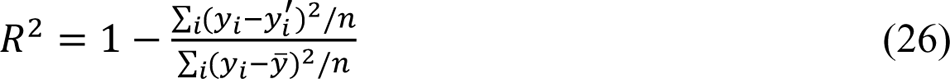

Where *y*_*i*_ and *y*^′^ represent the measured and simulated values, respectively; *y*^-^ is the mean value of measurement; n is the sample size.

## 3. Result

In this section, the response of DRF to canopy photosynthesis was analyzed in section 3.1. First, the study investigated the sensitivity of the DRF to APAR, which was validated by measured FAPAR. Next, the analysis looked at the sensitivity of the DRF to canopy photosynthesis, which was based on the validation of the APAR. The response of LCC to leaf and canopy photosynthesis was analyzed in section 3.2. Leaf A_max_ and α simulated by TPMSCC were validated, sensitivity analysis of canopy photosynthesis to LCC was conducted. The simulated accuracy of WheatGrow-T was analyzed in section 3.3; The capacity of WheatGrow-T, WheatGrow-B, and WheatGrow to capture the AGB time sequence under different N application rates was compared. The sensitivity of Cab_max_ to simulated agronomy parameters (AGB, LCC, LNC) was also analyzed. Finally, the simulated accuracy of agronomy parameters after the calibration of the WheatGrow-T and WheatGrow models were compared.

### 3.1 Canopy photosynthesis vs. diffuse radiation fraction

#### 3.1.1 Difference of APAR_dir_ and APAR_dif_

The comparison between the FAPAR simulation between TPMSCC and Big-leaf was conducted across the jointing, booting, and heading stages (Fig. 3). The results indicated that TPMSCC performed better than the Big-leaf model during all stages. TPMSCC mitigated the overestimation during the jointing stage and the underestimation during the booting and heading stages of the Big-leaf model (Fig. 3), which cued the relative magnitude between Big-leaf and TPMSCC varied in different environment and canopy structure. Figure 4 revealed the relative difference between FAPAR simulated by TPMSCC and Big-leaf, the absolute difference expended with the increase of DRF, and the relative size was highly depended on *θ_s_*. The incident light is composed by direct light and diffused light, but Big-leaf model treated all incident light as direct light, thus the variation of relative size revealed the relative size between APAR_dir_ and APAR_dif_ varied with environment and canopy structure. The sensitive analysis of TPMSCC model is necessary to explore the impact of environmental factors on direct and diffused light absorption to comprehend the advantage of TPMSCC.

**Fig. 3:**
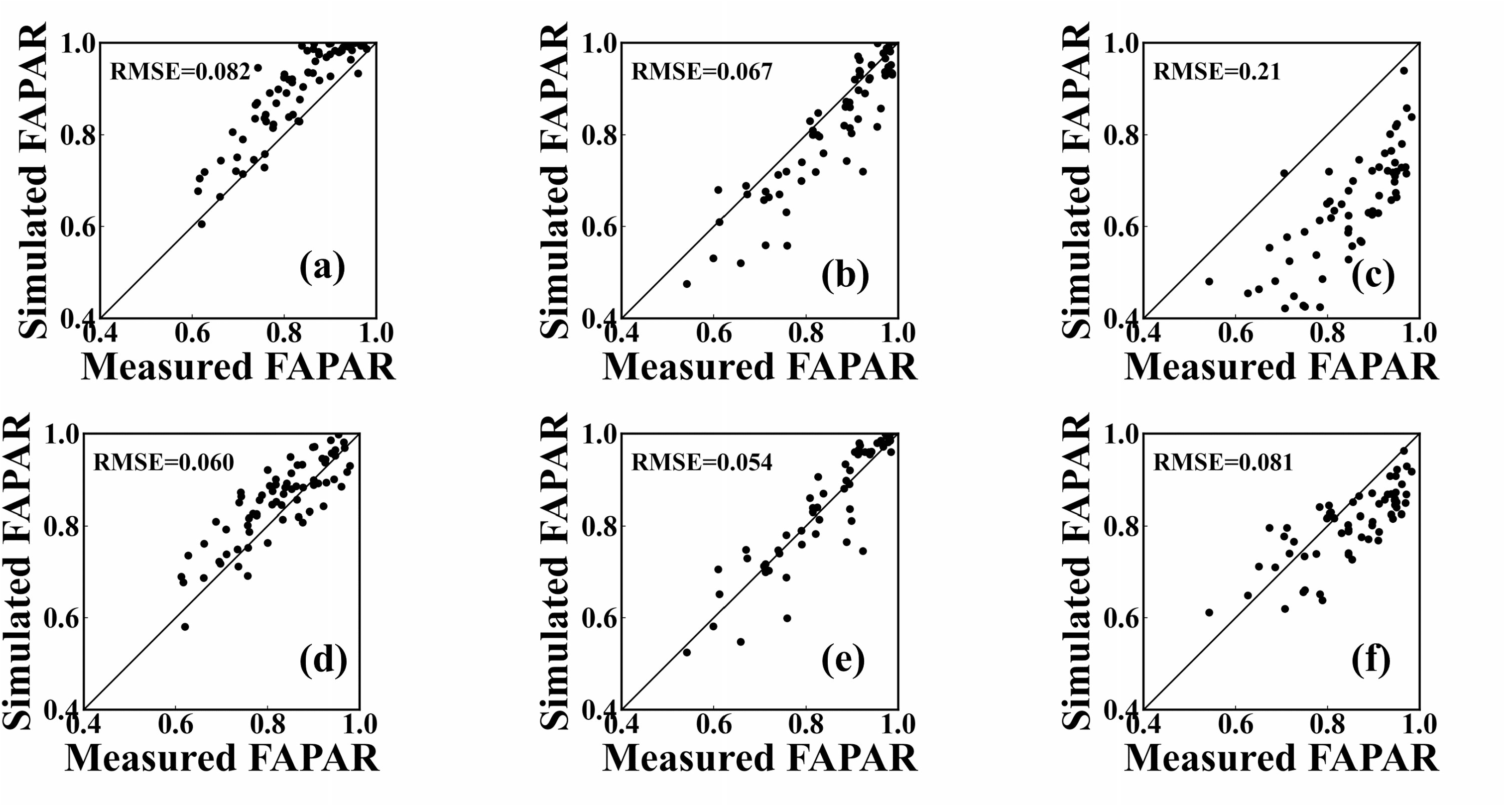
Comparison between the measured and simulated FAPAR values by TPMSCC and Big-leaf model. (a-c) show results simulated by the Big-leaf model; (d-f) display results simulated by TPMSCC. (a, d), (b, e), and (c, f) represent results at wheat jointing, booting, and heading stages.

**Fig. 4:**
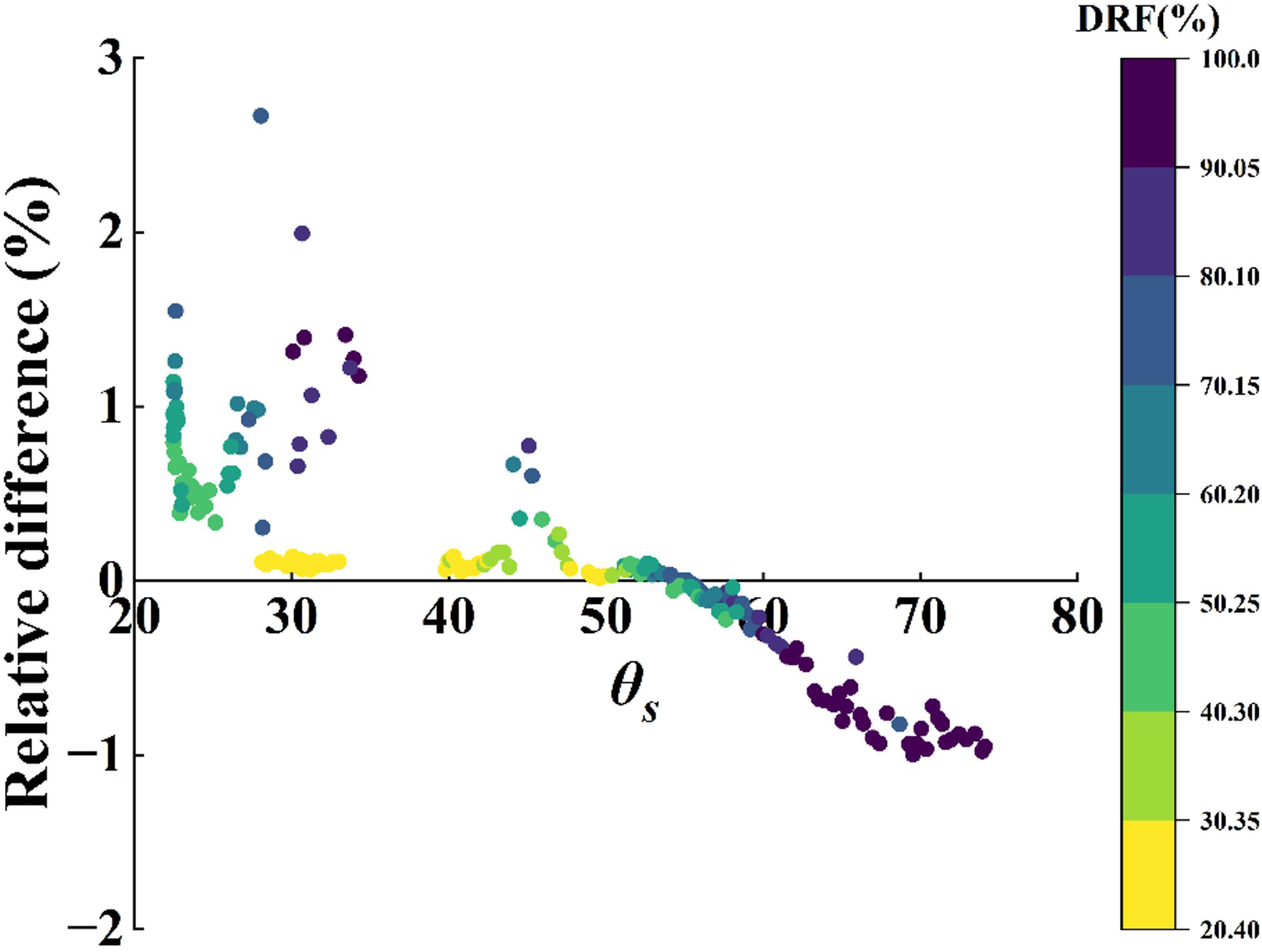
Relationship between the *θ_s_* and relative difference ([FAPAR_TPMSCC_-FAPAR_Big-leaf_]/FAPAR_Big-leaf_) in FAPAR simulated by TPMSCC and Big-leaf model.

The sensitivity analysis of APAR_dir_ and APAR_dif_ was conducted under different canopy densities and *θ_s_* (Fig. 5). When solar was at zenith (*θ_s_* = 0°), vertical distribution of APAR_dir_ was more uniform than APAR_dif_, but the vertical integral of APAR_dir_ was 47% (Fig. 5a) and 12% (Fig. 5c) lower than APAR_dif_, in sparse and dense canopy, respectively. When solar was near the horizon (*θ_s_* = 60°, the situation when solar was at the horizon was not analyzed because the absence of direct light during nightfall), the vertical distribution of APAR_dir_ and APAR_dif_ presented as a similarly skewed distribution, the vertical integral of APAR_dif_ was also similar with APAR_dir_ (just 7% higher and 4% lower than APAR_dir_ in sparse and dense canopy, respectively). When solar was at its zenith, the increase of canopy density increased APAR_dir_ with an increase of 38%, while the increase of APAR_dif_ was just 5% (Fig. 5a, c). When solar was near the horizon, the increase in canopy density resulted in only a 6% increase in APAR_dir_ and a 5% decrease in APAR_dif_ (Fig. 5b, d). The APAR_dir_ under high solar zenith angle raised by 17% and 52% in dense and sparse canopy, respectively, compared with that under low solar zenith angle, while the APAR_dif_ remained stable under varied solar zenith angle.

**Fig. 5:**
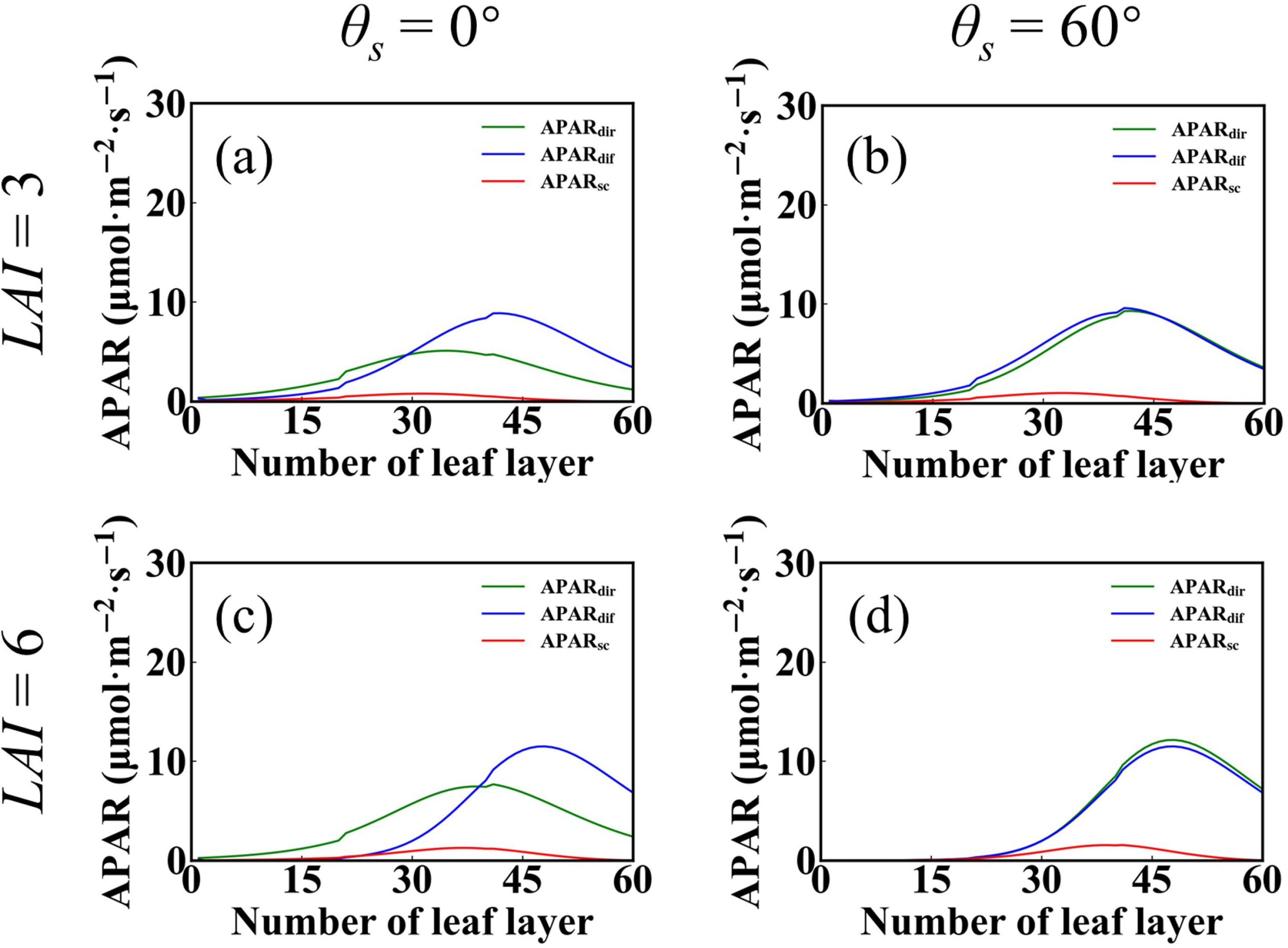
Absorbed Photosynthetically Active Radiation (APAR) of each leaf layer under different *θ_s_* and canopy density. (a, b) and (c, d) were the results of sparse canopy and dense canopy, respectively; (a, c) and (b, d) were the results of high *θ_s_* and low *θ_s_*, respectively. The DRF equaled 50%.

#### 3.1.2 Response of diffuse radiation fraction on photosynthesis

The sensitivity of A_canopy_ to PAR revealed that A_canopy_ increased first and then decreased (Fig. 6) in response to increasing PAR under natural illumination, cuing that the photosynthesis under sunny day is lower than that under slightly cloudy day. The decrease was more apparent when solar was near the horizon than at the zenith (Fig. 6b, d). A_canopy_ generally increased with rising DRF (Fig. 6a, b, d) under ideal illumination. However, for the dense canopy exposed to low θ_s_ values, A_canopy_ initially increased and then decreased with the increase in DRF (Fig. 6c).

**Fig. 6:**
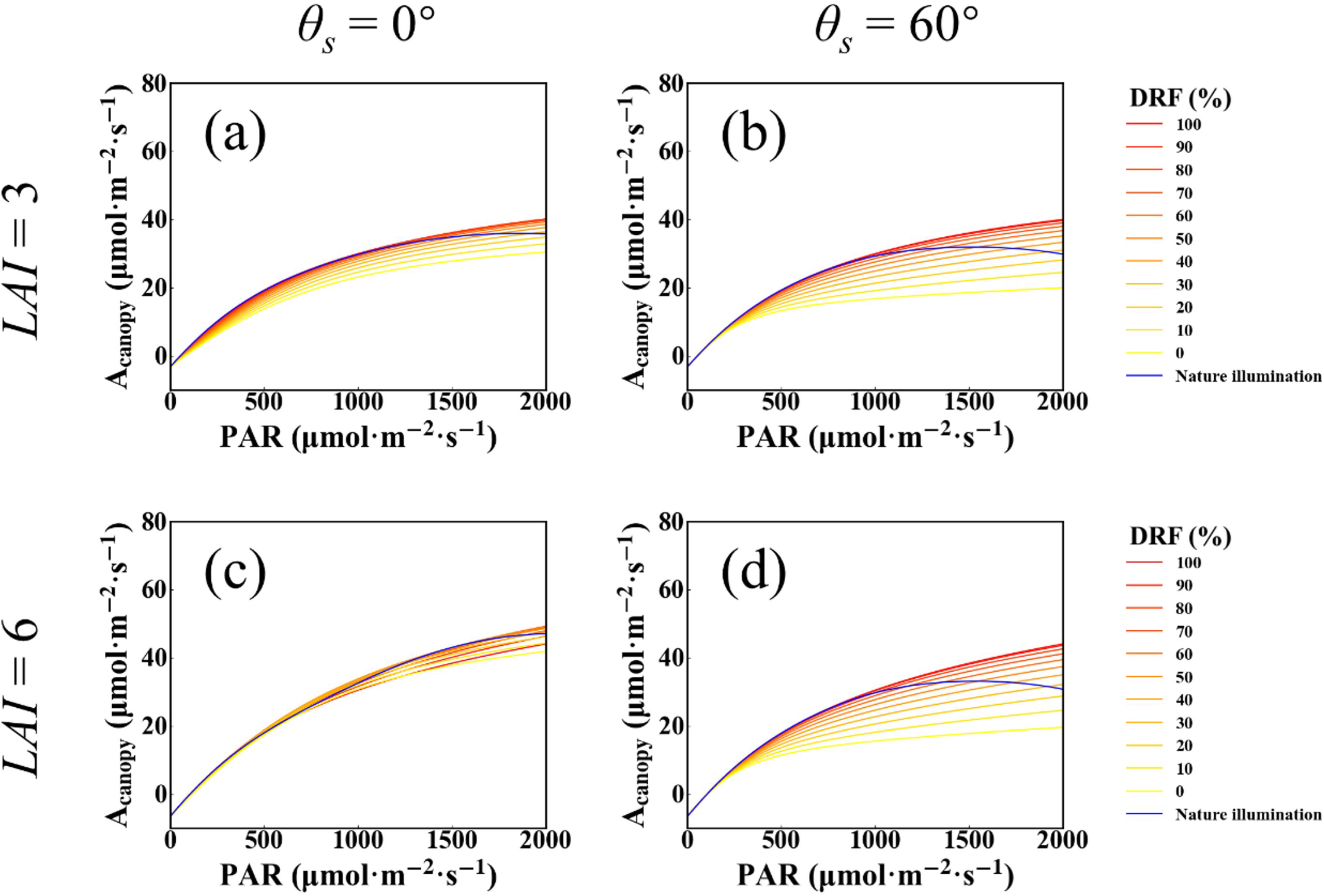
Light response curves simulated by TPMSCC under different DRFs (ideal illumination) and natural illumination when *θ_s_* equaled 0° (a, c) and 60° (b, d). Sparse canopy conditions are shown in (a, b), while dense canopy conditions are displayed in (c, d) (taking LAI 3 and 6 as examples).

The difference between A_canopy_ simulated by TPMSCC and Big-leaf_sep_ was apparent in high PAR value. The canopy light response curve simulated by TPMSCC had an obvious light saturation point (Fig. 6), while Big-leaf_sep_ failed to simulate this feature (Fig. 7).

**Fig. 7:**
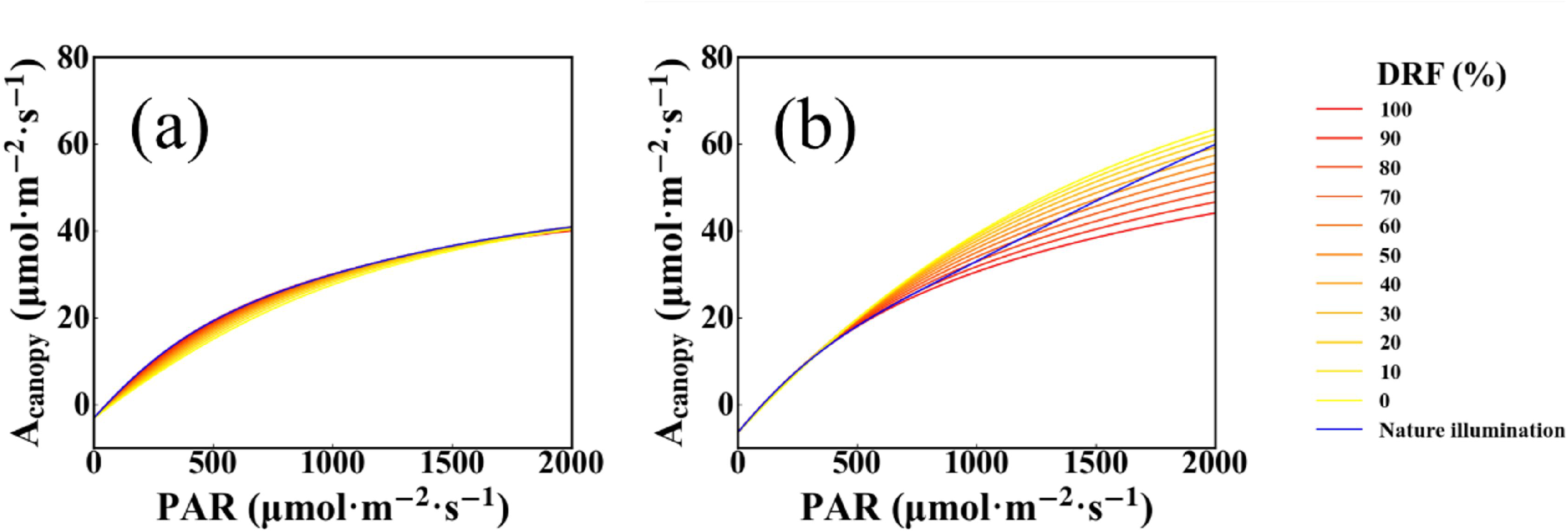
Light response curves simulated by Big-leaf_sep_ under different DRFs (ideal illumination) and natural illumination when *θ_s_* equaled 0°. Sparse canopy (a) and dense canopy (b) conditions are presented (taking LAI=3 and 6 as examples).

### 3.2 Leaf chlorophyll content vs. photosynthesis

#### 3.2.1 Effects on leaf photosynthesis

The trend of the leaf light response curve with varied LCC simulated by TPMSCC is shown in Figure S3. There was a high sensitivity of LCC to leaf photosynthetic rate, A_max_ and α were linearly and logarithmically related to LCC, respectively (Fig. 8). These simulated relationships corresponded closely with the results of Experiment 3. The R^2^ value between the measured and simulated A_max_ values was 0.39, RMSE was 3.66 μmol·m^-2^·s^-1^; R^2^ for simulated α was 0.42, RMSE was 0.0069.

**Fig. 8:**
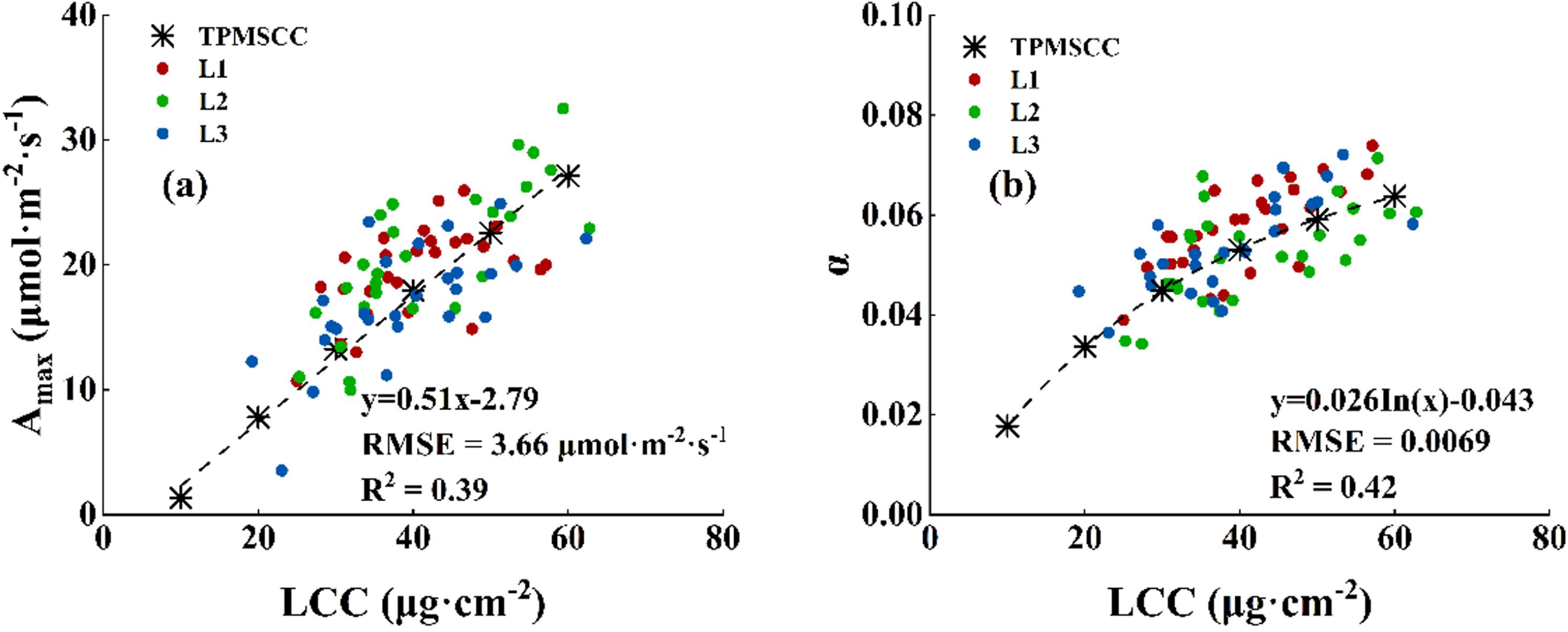
Relationships between photosynthetic parameters (A_max_ and α) and LCC. The asterisk represents results simulated by TPMSCC; L1, L2, and L3 were measured values of the first, second, and third fully expanded leaves from the top in Exp.3.

#### 3.2.2 Effect on Canopy Photosynthesis

There was a high sensitivity of LCC_Canopy_ to APAR, the APAR of canopy with high LCC_Canopy_ was 70% higher than that with low LCC_Canopy_ (Fig. S4). In addition, the proportions of APAR_sc_ to total APAR under high and low LCC_canopy_ were 6% and 12%, respectively (Fig. S4).

Canopy light response curves were simulated by the TPMSCC model at different LCC_canopy_ (Fig. S5), A_max,_ and α were extracted (Fig. S6). The correlations of A_max_ and α to LCC_canopy_ simulated by TPMSCC were good linear and logarithmic, respectively (Fig. S6). A_max_ and α of high LCC_canopy_ canopy were 6.3 and 2.3 times higher than that of low LCC_canopy_ canopy, respectively.

### 3.3 Evaluation of WheatGrow-T

#### 3.3.1 Effects of photosynthetic module

Three wheat growth models (WheatGrow-T, WheatGrow-B, and WheatGrow) were used to simulate AGB at different N levels, respectively. The findings demonstrated that WheatGrow-T accurately reflected the variations in AGB at various growth stages and N levels (Fig. 9a). WheatGrow-B captured the trend of AGB in different growth stages and N rates, but there was an obvious overestimation (Fig. 9b). However, WheatGrow could not simulate the variation trend of AGB with N rate and growth stage, and exhibited overestimation and underestimation at the beginning and end growth stages under low N treatment, respectively (Fig. 9c). The R^2^ values of AGB between the measured and simulated by WheatGrow-T were 0.84 and 0.89 for low and high N levels, respectively; while that for WheatGrow-B and WheatGrow were 0.73, 0.50, and 0.77, 0.72, respectively.

**Fig. 9:**
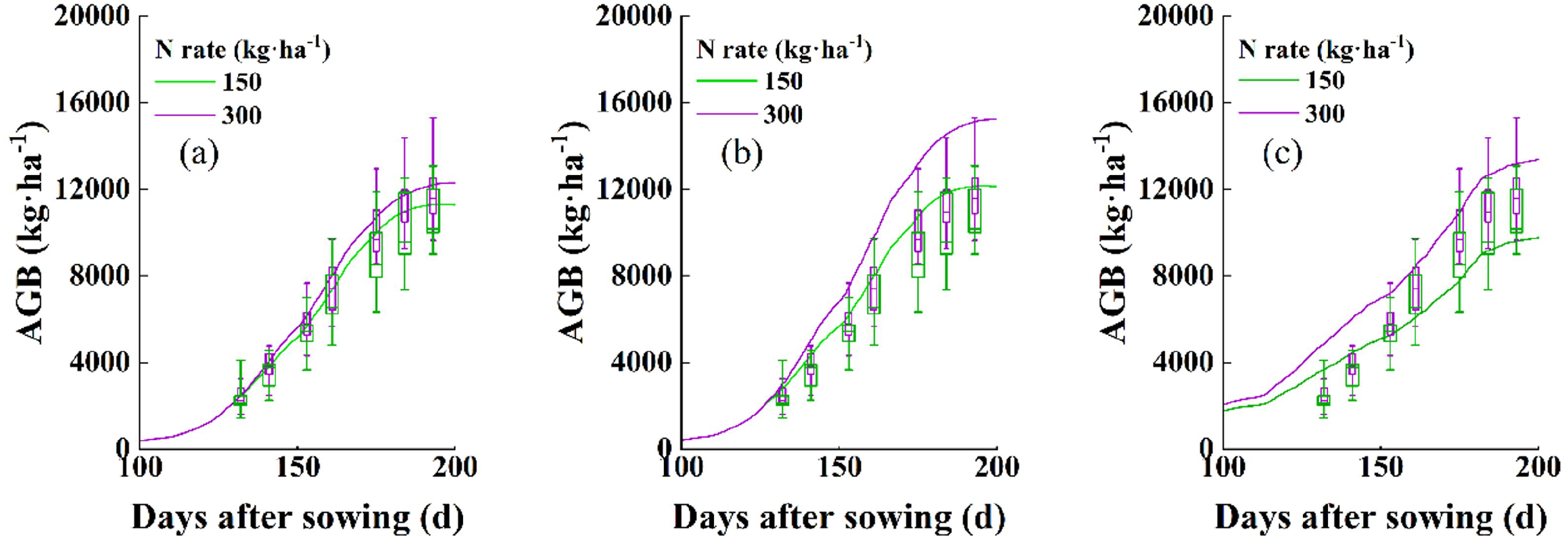
Box line plots comparing measured above-ground biomass (AGB) and simulated AGB by WheatGrow-T (a), WheatGrow-B (b), and WheatGrow (c) across different nitrogen application.

#### 3.3.2 Sensitivity of wheat growth parameters to N application

Two wheat varieties (NM22 and YM25) with wide differences in Cab_max_ were used as examples to analyze the differences in LNC, LCC_canopy_ and AGB under different N levels (Fig. 10). The measured maximum LCC values were 41 μg·cm^-2^ and 57 μg·cm^-2^ for NM22 and YM25, respectively. NM22 showed small differences between the high and low N levels, while YM25 showed large differences in LNC, LCC_canopy_, and AGB. In addition, NM22 had higher LNC, LCC_canopy,_ and AGB than YM25 under a low N application; while had lower LNC, LCC_canopy,_ and AGB than YM25 under a high N application.

**Fig. 10:**
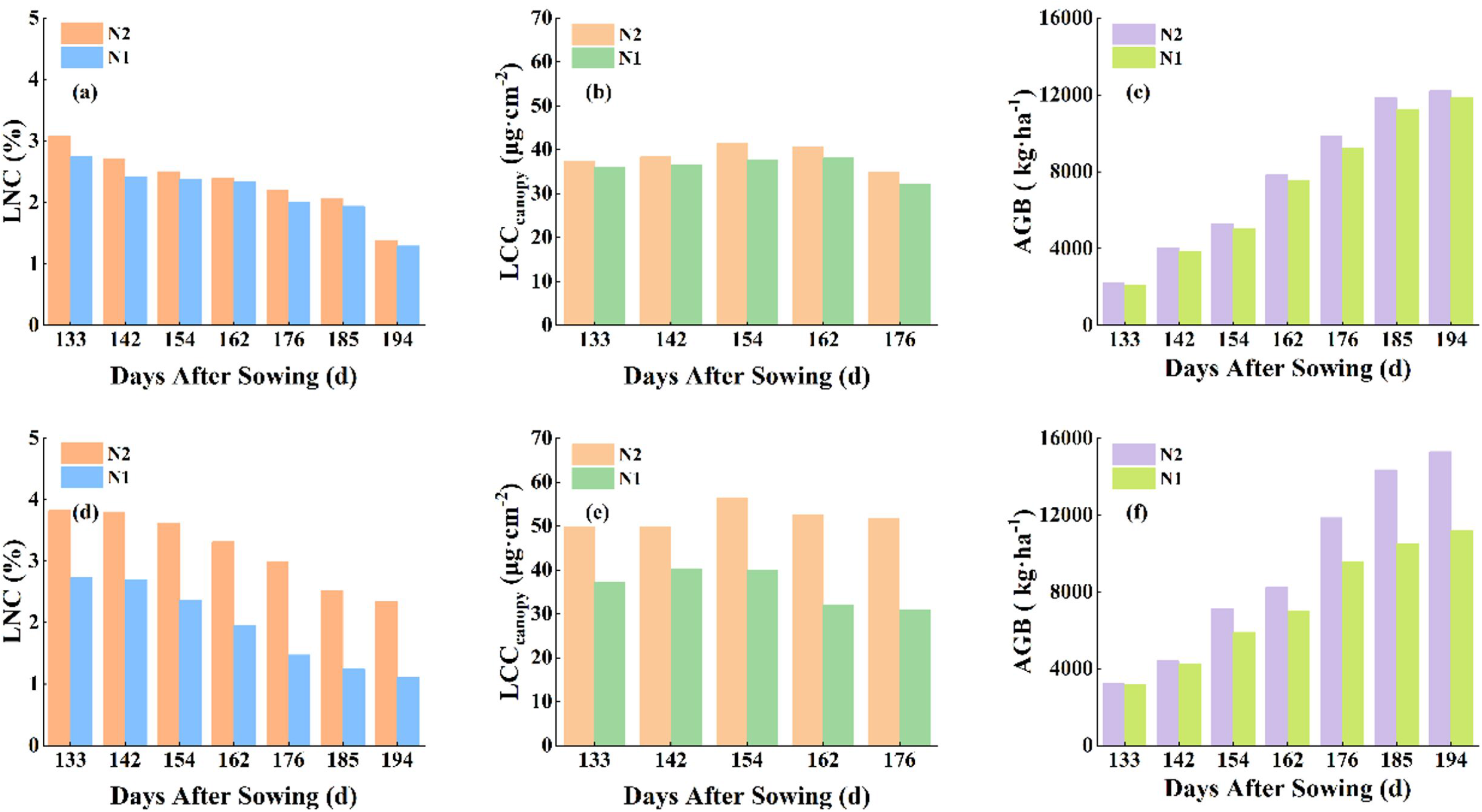
Measured leaf nitrogen content (LNC) (a, d), canopy leaf chlorophyll content (LCC_canopy_) (b, e), and above-ground biomass (AGB) (c, f) at various nitrogen application. (a b d) and (d e f) represent the results of NM22 and YM25, respectively.

The response of Cab_max_ to wheat growth parameters simulated by WheatGrow-T corresponded to the field measurements. Cab_max_ had significant effects on the simulated growth parameters of WheatGrow-T (Fig. 11). The simulation results also showed that wheat cultivar with high Cab_max_ had a higher sensitivity of LNC, LCC, and AGB to N level than that with low Cab_max_. The wheat cultivar with high Cab_max_ yielded higher LNC, LCC_canopy_, and AGB than that with low Cab_max_ at high N lever; however, the cultivar with high Cab_max_ was high in the vegetative stages and low in the reproductive stages at low N lever.

**Fig. 11:**
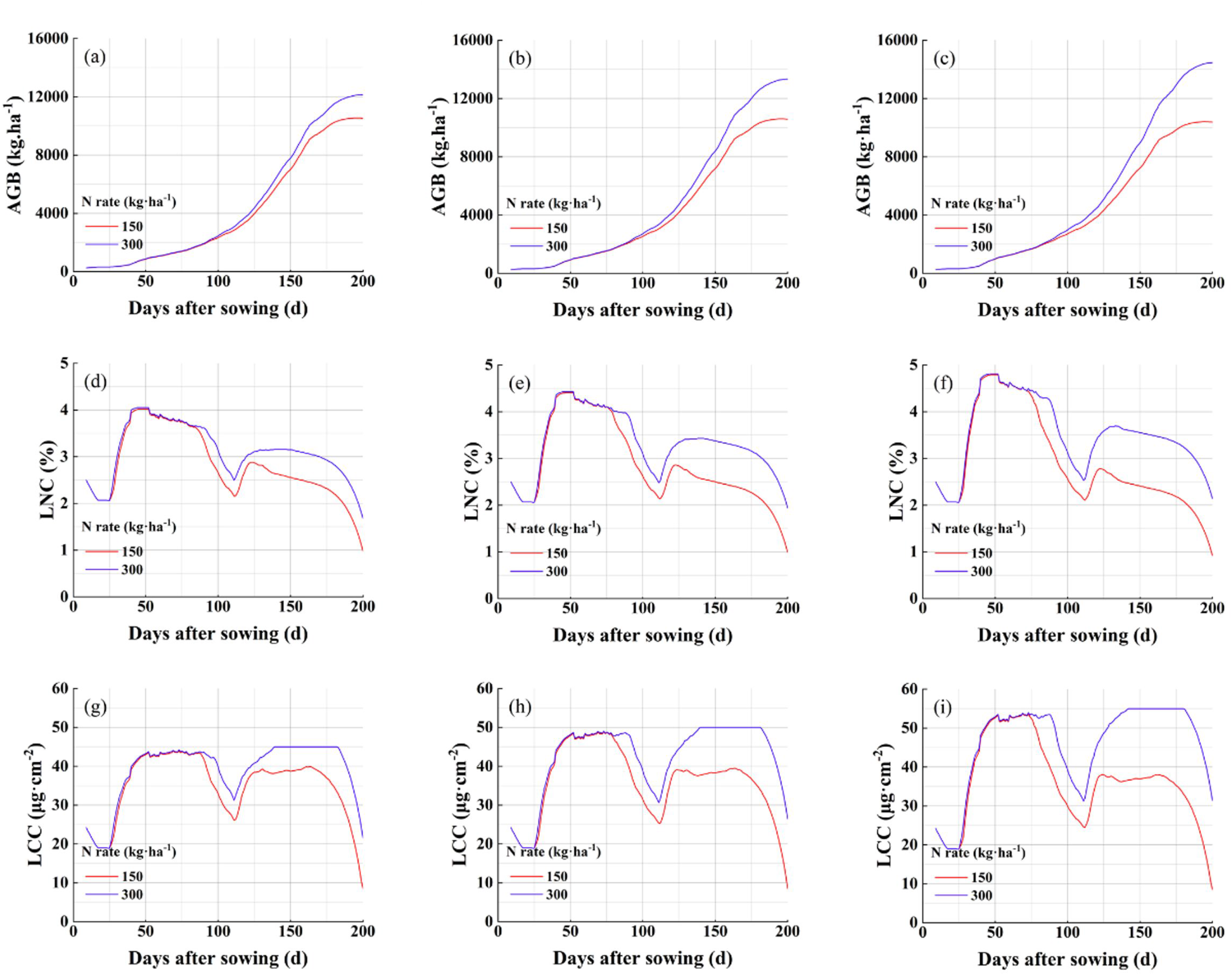
Time series of above-ground biomass (AGB) (a,b,c), leaf nitrogen content (LNC) (d,e,f), and leaf chlorophyll content (LCC) simulated by WheatGrow-T. The figures in first, second, and third columns correspond to wheat cultivars with Cab_max_ of 45 μg·cm^-2^, 50 μg·cm^-2^, and 55 μg·cm^-2^, respectively.

#### 3.3.3 Simulation accuracy of WheatGrow-T

AGB, LAI, LNC, LCC, and grain yield simulated by the WheatGrow-T model are shown in Figure 12. The results show that the simulated results of WheatGrow-T after calibration for Cab_max_ were better than those before calibration, which indicated that Cab_max_ was a key and reliable variety parameter. Before calibration of Cab_max_, the estimated results were overestimated in wheat varieties with low Cab_max_, whereas better performance was obtained for varieties with high Cab_max_. The overestimation was well resolved after the calibration of Cab_max_.

**Fig. 12:**
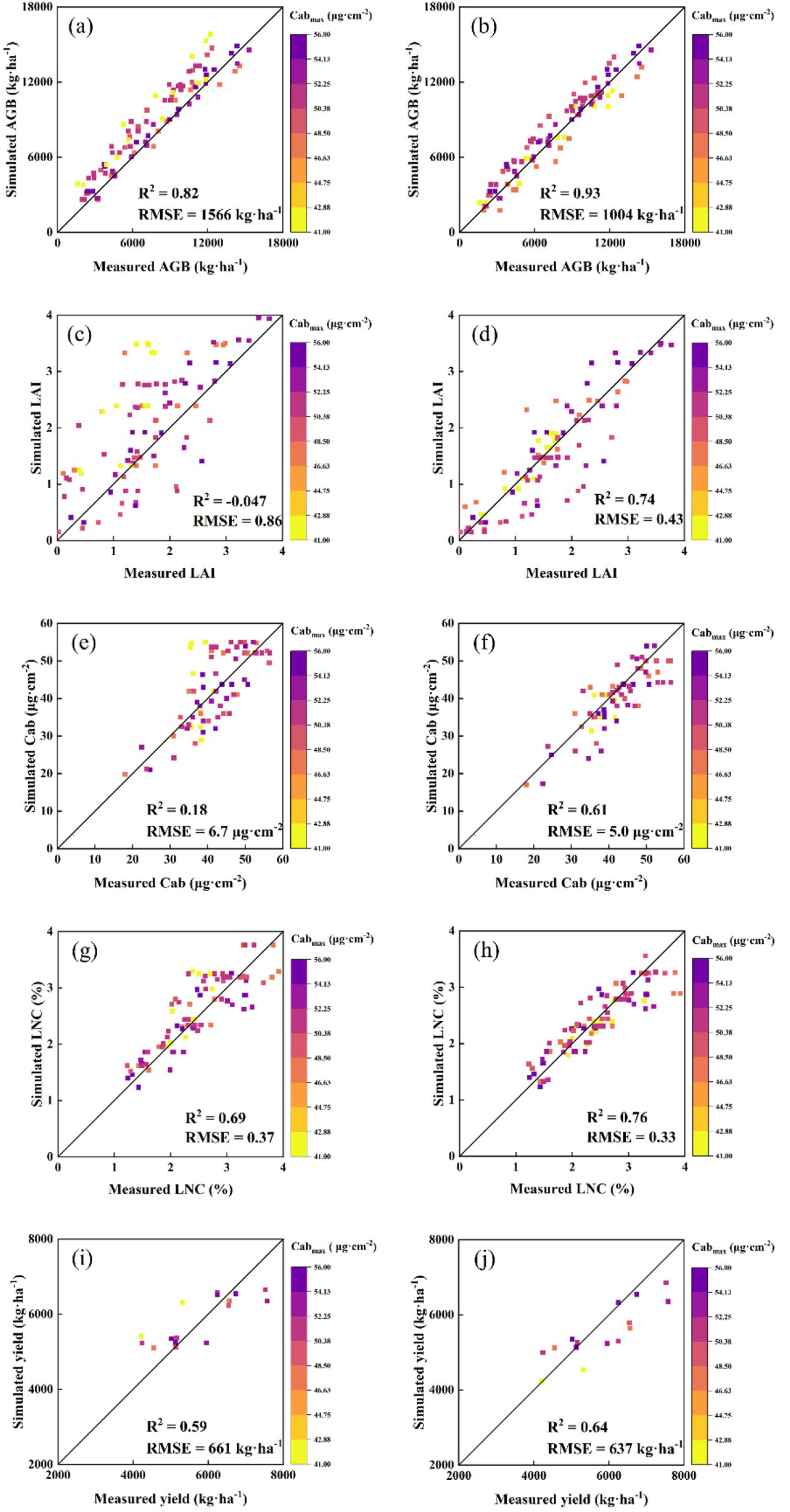
Comparison between simulated and measured values of WheatGrow-T. The figures in the first and second columns display simulated values before and after Cab_max_ calibration.

After gradually calibrating phenology, critical N concentration, and photosynthetic parameters, estimation accuracy of WheatGrow gradually increased. However, the estimation accuracy of WheatGrow was lower than WheatGrow-T under all the model calibration strategies (Table 6).

**Table 6:** Simulated accuracy of WheatGrow with calibration of various Parameters.

| Simulated value | Calibrated parameter |  |  |  |  |  |
| --- | --- | --- | --- | --- | --- | --- |
|  | Phenology |  | Phenology |  | Phenology |  |
|  |  |  | Critical nitrogen concentration |  | Critical nitrogen concentration |  |
|  |  |  |  |  | Photosynthetic parameter |  |
|  | RMSE | R<sup>2</sup> | RMSE | R<sup>2</sup> | RMSE | R<sup>2</sup> |
| AGB | 1673 kg·ha<sup>-1</sup> | 0.77 | 1589 kg·ha<sup>-1</sup> | 0.82 | 1241 kg·ha<sup>-1</sup> | 0.89 |
| LAI | 1.09 | -0.091 | 0.81 | 0.22 | 0.59 | 0.72 |
| LNC | 0.49 | 0.64 | 0.33 | 0.67 | 0.32 | 0.72 |
| Yield | 735 kg·ha<sup>-1</sup> | 0.41 | 708 kg·ha<sup>-1</sup> | 0.50 | 631 kg·ha<sup>-1</sup> | 0.55 |

## 4. Discussion

### 4.1 Response of diffuse radiation fraction to canopy photosynthesis

There was an opinion that diffuse radiation penetrates deeper into the crop canopy, so more uniformly distributed over all the leaves in the plant canopy compared with direct radiation (Kanniah et al., 2012; Li and Yang, 2015; Wang et al., 2018), which suggest the APAR_dir_ is higher than APAR_dif_. However, the finding that canopy has a higher FIPAR on overcast days than on clear days (Li and Fang, 2015; Shao et al., 2020) suggested the inconsistent result in diffuse light research. This study found the relative magnitude of APAR_dir_ and APAR_dif_ varied in different canopy structure and solar position, interpreting the contradiction in these studies. The FAPAR of direct light equals the projected area of leaf area at the direction of the solar incident, and increases with the rise of the solar zenith (Morris, 1989), the FAPAR of diffuse light can be treated as an integral of direct light interception at all the solar zenith (Monsi, 2004), thus APAR_dif_ could be higher than APAR_dir_ under low *θ_s_*. Result showed that the effect of canopy structure and solar position on APAR_dir_ and APAR_dif_ was different, APAR_dif_ was insensitive to the variation of canopy structure and solar position, APAR_dir_ showed higher sensitivity than APAR_dif_. The Big-leaf model treats all incident light as direct light, which means the difference in magnitude between the Big-leaf model and TPMSCC stems from the contrast between APAR_dif_ and APAR_dir_. The FAPAR simulated by Big-leaf model was similar with that by TPMSCC under the condition of low DRF, the higher accuracy of TPMSCC under illustration with high DRF demonstrates it well described the response of APAR to diffuse light.

Previous studies found that the canopy photosynthetic rate under slightly cloudy was higher than that under sunny days (Gu et al., 1999; Jing et al., 2010; Law et al., 2002; Oliveira et al., 2007; Yamasoe et al., 2006) because the diffuse light benefited the photosynthesis. When solar was near horizon, simulation results of TPMSCC well described the feature that highest photosynthetic rate occurred in slightly cloudy day (Fig. 6b, d), but such phenomenon was not apparent under low *θ_s_* (Fig. 6a, c). The variation of distribution characteristic and the total value of APAR under varied canopy density and solar position (Fig. 5) resulted in this variation. It is undoubtable that the horizontal distribution of APAR_dif_ was always more uniform than that of APAR_dir_ (Boote and Pickering, 1994; Yang et al., 2017). Meanwhile, the total APAR_dif_ was lower than APAR_dir_ when solar was at zenith, especially in sparse canopy. However, the vertical distribution of APAR_dir_ was more normal than that of APAR_dif_ when solar was at zenith (Fig. 5a, c). Thus, photosynthetic rate dropped when the advantage of diffuse light that uniform horizontal distribution and high total absorption can not make up the disadvantage of skewed vertical distribution. Therefore, when examining the impact of DRF on crop growth, it is necessary to take account the diurnal variations in solar position, light intensity, DRF, and canopy density.

Some scholars (Chen et al., 1999; De Pury and Farquhar, 1997) have found the overestimation of the big-leaf model on photosynthetic rate. An apparent overestimation by the big-leaf model was also found in the high intensity of PAR in this study (Fig. 6, Fig. 7). Additionally, it was clear that AGB had been overestimated (Fig. 9b). As a result, crop modeling’s photosynthetic module urgently needs to be improved. The overestimation of AGB was more serious in high N rate treatment than in low N rate treatment (Fig. 9b), which confirmed that the overestimation of photosynthetic rate was more pronounced in dense canopy compared to a sparse canopy (Fig. 6, Fig. 7). Canopy light response curve based on TPMSCC was corresponded to the field measurement by Song et al. (2022), and high simulating accuracy of AGB proved the reliability of TPMSCC model.

### 4.2 Response of leaf chlorophyll content on photosynthesis

The relationship of LCC to A_max_ and *ɑ* simulated by TPMSCC well interpreted the variation of field measurement (Fig. 8), and the linear and logarithms relationships were similar to the results of previous studies (Braune et al., 2009; Luo et al., 2019). Response of chlorophyll on photosynthesis was manifested in two aspects, one is light-harvesting pigments capture light energy and transmit it to reaction center pigments; the other is the conversion of light energy to electric energy, containing the motivation of reaction center pigments and the electron transport by electron transport chain (Jin et al., 2016). TPMSCC described these two processes by simulation of FAPAR_leaf_ and *J_max_*, which affected *ɑ* and A_max_, respectively. Previous research assumed that leaves either absorb all radiation (Zheng et al., 2015) or that the absorption fraction is constant (Liu et al., 2021). However, a high variation of FAPAR_leaf_ under varied LCC has been found in this study (Fig. S4). Yin et al. (2009) indicated that the impact of N on the conversion efficiency of incident light into J at the strictly limiting light level (k_2(LL)_) could be interpreted by the variation of leaf absorptance, which was simulated by the PROSPECT model in this study. As presented in equation (21), LCC affected FAPAR_leaf_, and then affected *ɑ*. The effect of N dynamic on A_max_ was determined by V_cmax_ or J_max_ (20, 22). Although the relationship between V_cmax_ and J_max_ varied with the environment, and a constant C value was hard to capture the variation of W_c_ to the stomatal conductance (Von Caemmerer et al., 2009), the relationship between LCC and A_max_ was well captured (Fig. 8a). In addition, several studies overlooked the diffuse light reflected by the background and leaves (Liu et al., 2021; Monsi, 2004). In this study, it was found that a proportion of 6% to 12% APAR came from the leaf diffuse light (Fig. S4), which indicated that the reabsorption of leaf scattered light should be considered.

Evans (1989) separated photosynthetic N into two parts, containing the soluble protein (RuBP carboxylase) and protein in the thylakoid membranes of the chloroplast, which corresponded to the dark reaction and light reaction, respectively. Chlorophyll and its protein complexes are the main participant of light reaction, which was the theoretical basis of the FvCB model to estimate *J_max_* using chlorophyll (Farquhar et al., 1980). Although chlorophyll is not involved in dark reaction directly, the plant optimizes N allocation to balance *V_c_* and *V_j_* (Chen et al., 1993; Evans, 1989; Wang et al., 2017), which results in a higher correlation between LCC and V_cmax_ than N concentration (Luo et al., 2019). These indicated that it is unreliable to estimate photosynthetic parameters using N dynamic directly. The reason may be that the allocation of photosynthetic N varied in different crop species, growth periods, and environmental conditions (Croft et al., 2017). It was more rational to estimate J_max_ than V_cmax_ using chlorophyll content (Croft et al., 2017; Luo et al., 2019). As a pigment with obvious absorption characteristics, chlorophyll could be easily estimated than soluble protein by the remote sensing (RS) method (Jiang et al., 2018; Li et al., 2020, 2018; Schlemmer et al., 2013). In addition, Evans (1989) found a relatively constant fraction of thylakoid protein under different illumination and N rates, but the fraction of soluble protein varied largely, so the correlation between chlorophyll and total N concentration is more stable.

### 4.3 Application of TPMSCC in crop models

The critical N concentration system cannot describe the difference in dry matter accumulation among wheat cultivars with different critical N concentrations without adjustment of photosynthetic parameters (Table 6). Although there were differences in photosynthetic parameters between cultivars (Harbinson and Yin, 2022), the variation might be explained by leaf physicochemical features (Fig. 8, Croft et al., 2017; Dechant et al., 2017). With no calibration of photosynthetic parameters, WheatGrow-T functioned well (Table 6). Evans (1989) found that the proportion of non-photosynthetic N increased as the total leaf N content increased, while the proportion of thylakoid proteins representing chlorophyll remained unchanged, which was the physiology basis of Cab_max_. The allocation of total leaf N to chlorophyll saturates as the leaf N content increases, and the increase of N was more composited as nucleic acids (Chapin and Kedrowski, 1983), thus increasing the respiration rate (Braune et al., 2009). The saturation of the electron transport rate and the continued increase of respiration rate with the increase of N content led to the existence of critical N concentration.

The adaptation of TPMSCC on crop models was well verified. The simulated accuracy of APAR and leaf photosynthetic parameters has been well validated, hence the calibration of WheatGrow-T did not depend on the measured AGB but still outperformed WheatGrow that calibrated by measured AGB. However, the variation of several components must be accounted when applying TPMSCC to other crop models and environments. Firstly, the variation of intercellular CO_2_ concentration affects W_c_ (17), thus affecting the relationship between A_max_ and N dynamic (Archontoulis et al., 2012). Although earlier research suggested that the RuBP-limited gross photosynthesis rate is the limiting factor for the carboxylation rate under atmospheric CO_2_ concentration (Evans, 1989; van der Tol et al., 2009), a free-air CO_2_ enrichment experiment suggested that the maximum rate of carboxylation also limited photosynthetic rate (Ainsworth and Long, 2005). A stomatal conductance model (Ball et al., 1987) will be added in the future to consider the variation of intercellular CO_2_ concentration. Secondly, the variation of the conversion coefficient from LNC to LCC also need to be considered, which varies under different environment and crops. The variation of the relationship between J_max_ and V_cmax_ also needs to be considered.

## 5. Conclusion

Traditionally, crop models have overlooked the impact of diffuse light in their light absorption simulations, and the representation of CO_2_ carboxylation rate lacked a robust physiological foundation. In response, this study coupled PROSAIL and FvCB to develop a novel photosynthetic model. PROSAIL was employed to simulate the I_abs_ and APAR. Simulated APAR is well-verified under different canopy densities and illumination conditions. Notably, our research revealed that the plant canopy significantly benefited from diffuse light. Subsequently, I_abs_ was input to FvCB to simulate the electron transfer rate and CO_2_ carboxylation rate. The simulated A_max_ and α had linear and logarithmic relationships to LCC because of the sensitivity of LCC on J_max_ and light absorption. These findings were validated through comprehensive field measurements. By coupling the TPMSCC and WheatGrow model, WheatGrow-T exhibited better performance on growth parameters and grain yield simulation than the original version (WheatGrow), even without calibrating for critical N concentrations and photosynthetic parameters. This strategy is a potential first step toward crop modeling that is more precise and physiologically grounded.

## Acknowledgments

This work was supported by the Science Fund for Creative Research Groups of the National Natural Science Foundation of China [grant number 32021004]; the National Key R&D Program of China [grant number 2022YFD2001102]; the National Natural Science Foundation of China [grant number 31971784]; the Jiangsu Collaborative Innovation Center for Modern Crop Production, the Priority Academic Program Development of Jiangsu Higher Education Institutions (PAPD); the China Postdoctoral Science Foundation [grant number 2022T150327]; and the National Science Fund for Distinguished Young Scholars [grant number 31725020].

## References

Alton, P.B., 2017. Retrieval of seasonal Rubisco-limited photosynthetic capacity at global FLUXNET sites from hyperspectral satellite remote sensing: Impact on carbon modelling. Agricultural and Forest Meteorology 232, 74–88. 10.1016/j.agrformet.2016.08.001

Archontoulis, S.V., Yin, X., Vos, J., Danalatos, N.G., Struik, P.C., 2012. Leaf photosynthesis and respiration of three bioenergy crops in relation to temperature and leaf nitrogen: how conserved are biochemical model parameters among crop species? Journal of Experimental Botany 63, 895–911. 10.1093/jxb/err321

Boote, K.J., Pickering, N., 1994. Modeling Photosynthesis of Row Crop Canopies. HortScience 29, 1423–1434. 10.21273/HORTSCI.29.12.1423

Braune, H., Müller, J., Diepenbrock, W., 2009. Integrating effects of leaf nitrogen, age, rank, and growth temperature into the photosynthesis-stomatal conductance model LEAFC3-N parameterised for barley (Hordeum vulgare L.). Ecological Modelling 220, 1599–1612. 10.1016/j.ecolmodel.2009.03.027

Chapin III, F.S., Kedrowski, R.A., 1983. Seasonal Changes in Nitrogen and Phosphorus Fractions and Autumn Retranslocation in Evergreen and Deciduous Taiga Trees. Ecology 64, 376–391. 10.2307/1937083

Chen, J.L., Reynolds, J.F., Harley, P.C., Tenhunen, J.D., 1993. Coordination theory of leaf nitrogen distribution in a canopy. Oecologia 93, 63–69. 10.1007/BF00321192

Chen, J.M., Liu, J., Cihlar, J., Goulden, M.L., 1999. Daily canopy photosynthesis model through temporal and spatial scaling for remote sensing applications. Ecological Modelling 124, 99–119. 10.1016/S0304-3800(99)00156-8

Cooper, M., Hammer, G.L., 1996. Synthesis of strategies for crop improvement. Plant adaptation and crop improvement. 591–623.

Cooper, M., Messina, C.D., Podlich, D., Totir, L.R., Baumgarten, A., Hausmann, N.J., Wright, D., Graham, G., 2014. Predicting the future of plant breeding: complementing empirical evaluation with genetic prediction. Crop Pasture Sci. 65, 311. 10.1071/CP14007

Croft, H., Chen, J.M., Luo, X., Bartlett, P., Chen, B., Staebler, R.M., 2017. Leaf chlorophyll content as a proxy for leaf photosynthetic capacity. Glob Change Biol 23, 3513–3524. 10.1111/gcb.13599

De Pury, D.G.G., Farquhar, G.D., 1997. Simple scaling of photosynthesis from leaves to canopies without the errors of big-leaf models. Plant Cell Environ 20, 537–557. 10.1111/j.1365-3040.1997.00094.x

de Wit, A., Boogaard, H., Fumagalli, D., Janssen, S., Knapen, R., van Kraalingen, D., Supit, I., van der Wijngaart, R., van Diepen, K., 2019. 25 years of the WOFOST cropping systems model. Agricultural Systems 168, 154–167. 10.1016/j.agsy.2018.06.018

Dechant, B., Cuntz, M., Vohland, M., Schulz, E., Doktor, D., 2017. Estimation of photosynthesis traits from leaf reflectance spectra: Correlation to nitrogen content as the dominant mechanism. Remote Sensing of Environment 196, 279–292. 10.1016/j.rse.2017.05.019

Dong, T., Liu, J., Shang, J., Qian, B., Ma, B., Kovacs, J.M., Walters, D., Jiao, X., Geng, X., Shi, Y., 2019. Assessment of red-edge vegetation indices for crop leaf area index estimation. Remote Sensing of Environment 222, 133–143. 10.1016/j.rse.2018.12.032

Dueri, S., Brown, H., Asseng, S., Ewert, F., Webber, H., George, M., Craigie, R., Guarin, J.R., Pequeno, D.N.L., Stella, T., Ahmed, M., Alderman, P.D., Basso, B., Berger, A.G., Mujica, G.B., Cammarano, D., Chen, Y., Dumont, B., Rezaei, E.E., Fereres, E., Ferrise, R., Gaiser, T., Gao, Y., Garcia-Vila, M., Gayler, S., Hochman, Z., Hoogenboom, G., Kersebaum, K.C., Nendel, C., Olesen, J.E., Padovan, G., Palosuo, T., Priesack, E., Pullens, J.W.M., Rodriguez, A., Roetter, R.P., Ramos, M.R., Semenov, M.A., Senapati, N., Siebert, S., Srivastava, A.K., Stockle, C., Supit, I., Tao, F., Thorburn, P., Wang, E., Weber, T.K.D., Xiao, L., Zhao, C., Zhao, J., Zhao, Z., Zhu, Y., Martre, P., 2022. Simulation of winter wheat response to variable sowing dates and densities in a high-yielding environment. J. Exp. Bot. 73, 5715–5729. 10.1093/jxb/erac221

Evans, J.R., 1989. Photosynthesis and nitrogen relationships in leaves of C3 plants. Oecologia 78, 9–19. 10.1007/BF00377192

Farquhar, G.D., von Caemmerer, S., Berry, J.A., 1980. A biochemical model of photosynthetic CO2 assimilation in leaves of C3 species. Planta 149, 78–90. 10.1007/BF00386231

Huang, H., Huang, J., Wu, Y., Zhuo, W., Song, J., Li, X., Li, L., Su, W., Ma, H., Liang, S., 2023. The Improved Winter Wheat Yield Estimation by Assimilating GLASS LAI Into a Crop Growth Model With the Proposed Bayesian Posterior-Based Ensemble Kalman Filter. IEEE Trans. Geosci. Remote Sensing 61, 1–18. 10.1109/TGRS.2023.3259742

Jacquemoud, S., Baret, F., 1990. PROSPECT: A model of leaf optical properties spectra. Remote Sensing of Environment 34, 75–91. 10.1016/0034-4257(90)90100-Z

Jay, S., Maupas, F., Bendoula, R., Gorretta, N., 2017. Retrieving LAI, chlorophyll and nitrogen contents in sugar beet crops from multi-angular optical remote sensing: Comparison of vegetation indices and PROSAIL inversion for field phenotyping. Field Crops Research 210, 33–46. 10.1016/j.fcr.2017.05.005

Jiang, C., Ryu, Y., 2016. Multi-scale evaluation of global gross primary productivity and evapotranspiration products derived from Breathing Earth System Simulator (BESS). Remote Sensing of Environment 186, 528–547. 10.1016/j.rse.2016.08.030

Jiang, J., Comar, A., Burger, P., Bancal, P., Weiss, M., Baret, F., 2018. Estimation of leaf traits from reflectance measurements: comparison between methods based on vegetation indices and several versions of the PROSPECT model. Plant Methods 14, 23. 10.1186/s13007-018-0291-x

Jiang, J., Weiss, M., Liu, S., Rochdi, N., Baret, F., 2020. Speeding up 3D radiative transfer simulations: A physically based metamodel of canopy reflectance dependency on wavelength, leaf biochemical composition and soil reflectance. Remote Sensing of Environment 237, 111614. 10.1016/j.rse.2019.111614

Jin, H., Li, M., Duan, S., Fu, M., Dong, X., Liu, B., Feng, D., Wang, J., Wang, H.-B., 2016. Optimization of Light-Harvesting Pigment Improves Photosynthetic Efficiency. Plant Physiol. 172, 1720–1731. 10.1104/pp.16.00698

Jing, X., Huang, J., Wang, G., Higuchi, K., Bi, J., Sun, Y., Yu, H., Wang, T., 2010. The effects of clouds and aerosols on net ecosystem CO_2_ exchange over semi-arid Loess Plateau of Northwest China. Atmospheric Chemistry and Physics 10, 8205–8218. 10.5194/acp-10-8205-2010

Jones, J.W., Hoogenboom, G., Porter, C.H., Boote, K.J., Batchelor, W.D., Hunt, L.A., Wilkens, P.W., Singh, U., Gijsman, A.J., Ritchie, J.T., 2003. The DSSAT cropping system model. European Journal of Agronomy 18, 235–265. 10.1016/S1161-0301(02)00107-7

Kang, Y., Özdoğan, M., 2019. Field-level crop yield mapping with Landsat using a hierarchical data assimilation approach. Remote Sensing of Environment 228, 144–163. 10.1016/j.rse.2019.04.005

Kanniah, K.D., Beringer, J., North, P., Hutley, L., 2012. Control of atmospheric particles on diffuse radiation and terrestrial plant productivity: A review. Progress in Physical Geography: Earth and Environment 36, 209–237. 10.1177/0309133311434244

Klosterhalfen, A., Herbst, M., Weihermüller, L., Graf, A., Schmidt, M., Stadler, A., Schneider, K., Subke, J.-A., Huisman, J.A., Vereecken, H., 2017. Multi-site calibration and validation of a net ecosystem carbon exchange model for croplands. Ecological Modelling 363, 137–156. 10.1016/j.ecolmodel.2017.07.028

Law, B.E., Falge, E., Gu, L., Baldocchi, D.D., Bakwin, P., Berbigier, P., Davis, K., Dolman, A.J., Falk, M., Fuentes, J.D., Goldstein, A., Granier, A., Grelle, A., Hollinger, D., Janssens, I.A., Jarvis, P., Jensen, N.O., Katul, G., Mahli, Y., Matteucci, G., Meyers, T., Monson, R., Munger, W., Oechel, W., Olson, R., Pilegaard, K., Paw U, K.T., Thorgeirsson, H., Valentini, R., Verma, S., Vesala, T., Wilson, K., Wofsy, S., 2002. Environmental controls over carbon dioxide and water vapor exchange of terrestrial vegetation. Agricultural and Forest Meteorology, FLUXNET 2000 Synthesis 113, 97–120. 10.1016/S0168-1923(02)00104-1

Lemaire, G., Salette, J., Sigogne, M., Terrasson, J.-P., 1984. Relation entre dynamique de croissance et dynamique de prélèvement d’azote pour un peuplement de graminées fourragères. I. — Etude de l’effet du milieu. Agronomie 4, 423–430. 10.1051/agro:19840503

Li, D., Chen, J.M., Zhang, X., Yan, Y., Zhu, J., Zheng, H., Zhou, K., Yao, X., Tian, Y., Zhu, Y., Cheng, T., Cao, W., 2020. Improved estimation of leaf chlorophyll content of row crops from canopy reflectance spectra through minimizing canopy structural effects and optimizing off-noon observation time. Remote Sensing of Environment 248, 111985. 10.1016/j.rse.2020.111985

Li, D., Cheng, T., Jia, M., Zhou, K., Lu, N., Yao, X., Tian, Y., Zhu, Y., Cao, W., 2018. PROCWT: Coupling PROSPECT with continuous wavelet transform to improve the retrieval of foliar chemistry from leaf bidirectional reflectance spectra. Remote Sensing of Environment 206, 1–14. 10.1016/j.rse.2017.12.013

Li, T., Yang, Q., 2015. Advantages of diffuse light for horticultural production and perspectives for further research. Frontiers in Plant Science 6.

Li, W., Fang, H., 2015. Estimation of direct, diffuse, and total FPARs from Landsat surface reflectance data and ground-based estimates over six FLUXNET sites. Journal of Geophysical Research: Biogeosciences 120, 96–112. 10.1002/2014JG002754

Liu, B., Martre, P., Ewert, F., Porter, J.R., Challinor, A.J., Müller, C., Ruane, A.C., Waha, K., Thorburn, P.J., Aggarwal, P.K., Ahmed, M., Balkovič, J., Basso, B., Biernath, C., Bindi, M., Cammarano, D., De Sanctis, G., Dumont, B., Espadafor, M., Eyshi Rezaei, E., Ferrise, R., Garcia-Vila, M., Gayler, S., Gao, Y., Horan, H., Hoogenboom, G., Izaurralde, R.C., Jones, C.D., Kassie, B.T., Kersebaum, K.C., Klein, C., Koehler, A.-K., Maiorano, A., Minoli, S., Montesino San Martin, M., Naresh Kumar, S., Nendel, C., O’Leary, G.J., Palosuo, T., Priesack, E., Ripoche, D., Rötter, R.P., Semenov, M.A., Stöckle, C., Streck, T., Supit, I., Tao, F., Van der Velde, M., Wallach, D., Wang, E., Webber, H., Wolf, J., Xiao, L., Zhang, Z., Zhao, Z., Zhu, Y., Asseng, S., 2019. Global wheat production with 1.5 and 2.0°C above pre-industrial warming. Global Change Biology 25, 1428–1444. 10.1111/gcb.14542

Liu, S., Baret, F., Abichou, M., Manceau, L., Andrieu, B., Weiss, M., Martre, P., 2021. Importance of the description of light interception in crop growth models. Plant Physiology 186, 977–997. 10.1093/plphys/kiab113

Luo, X., Croft, H., Chen, J.M., Bartlett, P., Staebler, R., Froelich, N., 2018. Incorporating leaf chlorophyll content into a two-leaf terrestrial biosphere model for estimating carbon and water fluxes at a forest site. Agricultural and Forest Meteorology 248, 156–168. 10.1016/j.agrformet.2017.09.012

Luo, X., Croft, H., Chen, J.M., He, L., Keenan, T.F., 2019. Improved estimates of global terrestrial photosynthesis using information on leaf chlorophyll content. Global Change Biology 25, 2499–2514. 10.1111/gcb.14624

Makowski, D., Zhao, B., Ata-Ul-Karim, S.T., Lemaire, G., 2020. Analyzing uncertainty in critical nitrogen dilution curves. European Journal of Agronomy 118, 126076. 10.1016/j.eja.2020.126076

Manschadi, A.M., Palka, M., Fuchs, W., Neubauer, T., Eitzinger, J., Oberforster, M., Soltani, A., 2022. Performance of the SSM-iCrop model for predicting growth and nitrogen dynamics in winter wheat. European Journal of Agronomy 135, 126487. 10.1016/j.eja.2022.126487

Messina, C.D., Podlich, D., Dong, Z., Samples, M., Cooper, M., 2011. Yield–trait performance landscapes: from theory to application in breeding maize for drought tolerance. Journal of Experimental Botany 62, 855–868. 10.1093/jxb/erq329

Monsi, M., 2004. On the Factor Light in Plant Communities and its Importance for Matter Production. Annals of Botany 95, 549–567. 10.1093/aob/mci052

Morris, J.T., 1989. Modelling light distribution within the canopy of the marsh grass Spartina alterniflora as a function of canopy biomass and solar angle. Agricultural and Forest Meteorology 46, 349–361. 10.1016/0168-1923(89)90036-1

Nelson, R.A., Holzworth, D.P., Hammer, G.L., Hayman, P.T., 2002. Infusing the use of seasonal climate forecasting into crop management practice in North East Australia using discussion support software. Agricultural Systems 74, 393–414. 10.1016/S0308-521X(02)00047-1

Oliveira, P.H.F., Artaxo, P., Pires, C., De Lucca, S., ProcóPio, A., Holben, B., Schafer, J., Cardoso, L.F., Wofsy, S.C., Rocha, H.R., 2007. The effects of biomass burning aerosols and clouds on the CO2 flux in Amazonia. Tellus B: Chemical and Physical Meteorology 59, 338–349. 10.1111/j.1600-0889.2007.00270.x

Parent, B., Tardieu, F., 2014. Can current crop models be used in the phenotyping era for predicting the genetic variability of yield of plants subjected to drought or high temperature? Journal of Experimental Botany 65, 6179–6189. 10.1093/jxb/eru223

Prioul, J.L., Chartier, P., 1977. Partitioning of Transfer and Carboxylation Components of Intracellular Resistance to Photosynthetic CO2 Fixation: A Critical Analysis of the Methods Used. Annals of Botany 41, 789–800. 10.1093/oxfordjournals.aob.a085354

Puntel, L.A., Sawyer, J.E., Barker, D.W., Dietzel, R., Poffenbarger, H., Castellano, M.J., Moore, K.J., Thorburn, P., Archontoulis, S.V., 2016. Modeling long-term corn yield response to nitrogen rate and crop rotation. Frontiers in Plant Science 7.

Gu, L., Fuentes, J.D., Shugart, H.H., Staebler, R.M., Black, T.A., 1999. Responses of net ecosystem exchanges of carbon dioxide to changes in cloudiness: Results from two North American deciduous forests. Journal of Geophysical Research: Atmospheres 104, 31421–31434. 10.1029/1999JD901068

Ryu, Y., Baldocchi, D.D., Kobayashi, H., van Ingen, C., Li, J., Black, T.A., Beringer, J., van Gorsel, E., Knohl, A., Law, B.E., Roupsard, O., 2011. Integration of MODIS land and atmosphere products with a coupled-process model to estimate gross primary productivity and evapotranspiration from 1 km to global scales. Global Biogeochemical Cycles 25. 10.1029/2011GB004053

Schlemmer, M., Gitelson, A., Schepers, J., Ferguson, R., Peng, Y., Shanahan, J., Rundquist, D., 2013. Remote estimation of nitrogen and chlorophyll contents in maize at leaf and canopy levels. International Journal of Applied Earth Observation and Geoinformation 25, 47–54. 10.1016/j.jag.2013.04.003

Shao, L., Li, G., Zhao, Q., Li, Y., Sun, Y., Wang, W., Cai, C., Chen, W., Liu, R., Luo, W., Yin, X., Lee, X., 2020. The fertilization effect of global dimming on crop yields is not attributed to an improved light interception. Global Change Biology 26, 1697–1713. 10.1111/gcb.14822

Song, Q., Van Rie, J., Den Boer, B., Galle, A., Zhao, H., Chang, T., He, Z., Zhu, X.-G., 2022. Diurnal and Seasonal Variations of Photosynthetic Energy Conversion Efficiency of Field Grown Wheat. Front. Plant Sci. 13, 817654. 10.3389/fpls.2022.817654

Soufizadeh, S., Munaro, E., McLean, G., Massignam, A., van Oosterom, E.J., Chapman, S.C., Messina, C., Cooper, M., Hammer, G.L., 2018. Modelling the nitrogen dynamics of maize crops – Enhancing the APSIM maize model. European Journal of Agronomy 100, 118–131. 10.1016/j.eja.2017.12.007

Spitters, C.J.T., Toussaint, H.A.J.M., Goudriaan, J., 1986. Separating the diffuse and direct component of global radiation and its implications for modeling canopy photosynthesis Part I. Components of incoming radiation. Agricultural and Forest Meteorology 38, 217–229. 10.1016/0168-1923(86)90060-2

Tang, Y., Zhou, R., He, P., Yu, M., Zheng, H., Yao, X., Cheng, T., Zhu, Y., Cao, W., Tian, Y., 2023. Estimating wheat grain yield by assimilating phenology and LAI with the WheatGrow model based on theoretical uncertainty of remotely sensed observation. Agricultural and Forest Meteorology 339, 109574. 10.1016/j.agrformet.2023.109574

Verhoef, W., 1984. Light scattering by leaf layers with application to canopy reflectance modeling: The SAIL model. Remote Sensing of Environment 16, 125–141. 10.1016/0034-4257(84)90057-9

Verrelst, J., Rivera, J.P., Veroustraete, F., Muñoz-Marí, J., Clevers, J.G.P.W., Camps-Valls, G., Moreno, J., 2015. Experimental Sentinel-2 LAI estimation using parametric, non-parametric and physical retrieval methods – A comparison. ISPRS Journal of Photogrammetry and Remote Sensing 108, 260–272. 10.1016/j.isprsjprs.2015.04.013

Von Caemmerer, S., Farquhar, G., Berry, J., 2009. Biochemical Model of C3 Photosynthesis, in: Laisk, A., Nedbal, L., Govindjee (Eds.), Photosynthesis in Silico, Advances in Photosynthesis and Respiration. Springer Netherlands, Dordrecht, pp. 209–230. 10.1007/978-1-4020-9237-4_9

Wang, H., Prentice, I.C., Keenan, T.F., Davis, T.W., Wright, I.J., Cornwell, W.K., Evans, B.J., Peng, C., 2017. Towards a universal model for carbon dioxide uptake by plants. Nature Plants 3, 734–741. 10.1038/s41477-017-0006-8

Wang, X., Wu, J., Chen, M., Xu, X., Wang, Z., Wang, B., Wang, C., Piao, S., Lin, W., Miao, G., Deng, M., Qiao, C., Wang, J., Xu, S., Liu, L., 2018. Field evidences for the positive effects of aerosols on tree growth. Glob Change Biol 24, 4983–4992. 10.1111/gcb.14339

Widlowski, J.L., 2010. On the bias of instantaneous FAPAR estimates in open-canopy forests. Agricultural and Forest Meteorology 150, 1501–1522. 10.1016/j.agrformet.2010.07.011

Wullschleger, S.D., 1993. Biochemical Limitations to Carbon Assimilation in C3 Plants—A Retrospective Analysis of the A/Ci Curves from 109 Species. Journal of Experimental Botany 44, 907–920. 10.1093/jxb/44.5.907

Xi, M., Qi, Z., Zou, Y., Raghavan, G.S.V., Sun, J., 2015. Calibrating RZWQM2 model using quantum-behaved particle swarm optimization algorithm. Computers and Electronics in Agriculture 113, 72–80. 10.1016/j.compag.2015.02.002

Xu, X.Q., Lu, J.S., Zhang, N., Yang, T.C., He, J.Y., Yao, X., Cheng, T., Zhu, Y., Cao, W.X., Tian, Y.C., 2019. Inversion of rice canopy chlorophyll content and leaf area index based on coupling of radiative transfer and Bayesian network models. ISPRS Journal of Photogrammetry and Remote Sensing 150, 185–196. 10.1016/j.isprsjprs.2019.02.013

Yamasoe, M.A., von Randow, C., Manzi, A.O., Schafer, J.S., Eck, T.F., Holben, B.N., 2006. Effect of smoke and clouds on the transmissivity of photosynthetically active radiation inside the canopy. Atmospheric Chemistry and Physics 6, 1645–1656. 10.5194/acp-6-1645-2006

YanDa, L., Liang, T., YuPing, Z., XiangCheng, Z., WeiXing, C., Yan, Z., 2010. Relationship of PAR interception of canopy to leaf area and yield in rice. Scientia Agricultura Sinica 43, 3296–3305.

Yang, P., Verhoef, W., van der Tol, C., 2017. The mSCOPE model: A simple adaptation to the SCOPE model to describe reflectance, fluorescence and photosynthesis of vertically heterogeneous canopies. Remote Sensing of Environment 201, 1–11. 10.1016/j.rse.2017.08.029

Ye, Z., Qiu, X., Chen, J., Cammarano, D., Ge, Z., Ruane, A.C., Liu, L., Tang, L., Cao, W., Liu, B., Zhu, Y., 2020. Impacts of 1.5 °C and 2.0 °C global warming above pre-industrial on potential winter wheat production of China. European Journal of Agronomy 120, 126149. 10.1016/j.eja.2020.126149

Yin, X., Harbinson, J., Struik, P.C., 2006. Mathematical review of literature to assess alternative electron transports and interphotosystem excitation partitioning of steady-state C3 photosynthesis under limiting light. Plant, Cell & Environment 29, 1771–1782. 10.1111/j.1365-3040.2006.01554.x

Yin, X., Struik, P.C., Romero, P., Harbinson, J., Evers, J.B., Van Der Putten, P.E.L., Vos, J., 2009. Using combined measurements of gas exchange and chlorophyll fluorescence to estimate parameters of a biochemical C3 photosynthesis model: a critical appraisal and a new integrated approach applied to leaves in a wheat (Triticum aestivum) canopy. Plant, Cell & Environment 32, 448–464. 10.1111/j.1365-3040.2009.01934.x

Zhang, N., Su, X., Zhang, X., Yao, X., Cheng, T., Zhu, Y., Cao, W., Tian, Y., 2020. Monitoring daily variation of leaf layer photosynthesis in rice using UAV-based multi-spectral imagery and a light response curve model. Agricultural and Forest Meteorology 291, 108098. 10.1016/j.agrformet.2020.108098

Zhao, C., Li, H., Li, P., Yang, G., Gu, X., Lan, Y., 2017. Effect of Vertical Distribution of Crop Structure and Biochemical Parameters of Winter Wheat on Canopy Reflectance Characteristics and Spectral Indices. IEEE Trans. Geosci. Remote Sensing 55, 236–247. 10.1109/TGRS.2016.2604492

Zhao, Z., Wang, E., Wang, Z., Zang, H., Liu, Y., Angus, J.F., 2014. A reappraisal of the critical nitrogen concentration of wheat and its implications on crop modeling. Field Crops Research 164, 65–73. 10.1016/j.fcr.2014.05.004

Zhou, K., Guo, Y., Geng, Y., Zhu, Y., Cao, W., Tian, Y., 2014. Development of a Novel Bidirectional Canopy Reflectance Model for Row-Planted Rice and Wheat. Remote Sensing 6, 7632–7659. 10.3390/rs6087632

